# Structure and Flexibility of the Yeast NuA4 Histone Acetyltransferase Complex

**DOI:** 10.1101/2022.06.24.497536

**Authors:** Stefan A. Zukin, Matthew R. Marunde, Irina K. Popova, Eva Nogales, Avinash B. Patel

## Abstract

The NuA4 protein complex acetylates histones H4 and H2A to activate both transcription and DNA repair. We report the 3.0 Å-resolution cryo-electron microscopy structure of the central hub of NuA4, which flexibly tethers the HAT and TINTIN modules. The hub contains the large Tra1 subunit and a core that includes Swc4, Arp4, Act1, Eaf1 and the C-terminal region of Epl1. Eaf1 stands out as the primary scaffolding factor that interacts with the Tra1, Sw4 and Epl1 subunits and contributes the conserved HSA helix to the Arp module. Using nucleosome binding assays, we find that the HAT module, which is anchored to the core through Epl1, recognizes H3K4me3 nucleosomes with hyperacetylated H3 tails, while the TINTIN module, anchored to the core via Eaf1, recognizes nucleosomes that have hyperacetylated H2A and H4 tails. Together with the known interaction of Tra1 with site-specific transcription factors, our data suggests a model in which Tra1 recruits NuA4 to specific genomic sites then allowing the flexible HAT and TINTIN modules to select nearby nucleosomes for acetylation.

## Introduction

Chemical modifications of histone proteins are a key mechanism by which gene expression is regulated. These chemical modifications can affect the physical state of chromatin, regulating whether it is in a tightly packed and repressed state, or in an open and active state^1–4^. One such chemical modification, lysine acetylation, is catalyzed by histone acetyltransferases (HATs), which are often part of large, multi-subunit complexes^5,6^. Acetylated histones can directly affect the stability of a nucleosome by neutralizing the otherwise positively charged lysines tails that help stabilize the binding of the octamer core to the negatively charged DNA, or indirectly by recruiting chromatin remodelers that can alter how histone octamers bind DNA^7–10^.

NuA4 (<u>Nu</u>cleosome <u>A</u>cetyltransferase of <u>H4</u>) is one of eight histone acetyltransferase (HAT)-containing complexes in *Saccharomyces cerevisiae*, and its catalytic subunit, Esa1 (or Kat5), is the only essential HAT in *S. cerevisiae*^11–20^. NuA4 is composed of a total of 13 subunits, which together give the complex a molecular weight of 1.3 MDa^16^. The complex is thought to be organized in four main parts: the Tra1 subunit, the core module, the HAT module and the TINTIN module^21^. Tra1, the largest component of NuA4, is a member of the phosphoinositide-3-kinase (PI3K)-related pseudo kinase (ΨPIKK) family of proteins, which lacks kinase activity and instead functions as the primary target for the binding of sequence specific transcription factors^22–25^. The core module has been proposed to include Eaf1, Swc4, Yaf9, Arp4 and Act1, and is thought to connect the other three parts of the complex together^16,21,25 26–28^. The HAT module (also known as Piccolo) contains Esa1, Yng2, Eaf6 and Epl1 and is responsible for the acetylation of histone H2A and H4 in target nucleosomes^16^. And lastly the TINTIN (Trimer Independent of NuA4 involved in Transcription Interactions with Nucleosomes) module is composed of Eaf3, Eaf5, and Eaf7, and has several speculated functions including binding the Pol II CTD, RNA and histones^29^.

To date there has only been one other cryo-electron microscopy (cryo-EM) structure of NuA4^28^. From this structure it was proposed that the Tra1 subunit and core module form a rigid connection, and the HAT module was only stably defined in a small subpopulation of the particles. However, mapping of the crosslinking mass spectrometry data published after the structure, showed that much of the de novo built regions of the structure were incompatible with the mass spectrometry data^28^. Additionally, the HAT module location was also described to be in a different location to later negative stain studies^28,30^.

Here we have used cryo-EM to visualize NuA4 and have resolved the stable central hub containing the Tra1 subunit and the core module at ∼3Å resolution, allowing for assignments of all the density within this region. We found that the core is composed of Eaf1, Swc4, Arp4 and Act1 as well as the C-terminus of Epl1. Based on our structure, we were able show that the flexible HAT and TINITIN modules are respectively tethered to the core through Epl1 and Eaf1. Using nucleosome binding assays, we were able to show that the HAT module prefers nucleosomes modified with H3K4me3 and hyperacetylated H3 tails (acetylation marks that are produced by the SAGA complex), while the TINTIN module has a weak preference for nucleosomes that are hyperacetylated at their H2A and H4 tails (the product of the NuA4 HAT module). Based on our findings, we propose a model of how NuA4 and other HAT complexes target nucleosomes for acetylation.

## RESULTS AND DISCUSSION

### Overall architecture of NuA4

For our structural studies we purified NuA4 from *S. cerevisiae* harboring a DNA fragment encoding a TAP tag at the 3’-end of the Esa1 acetyl transferase gene. The isolated NuA4 contained all 13 subunits of the complex (Fig. 1A), as confirmed by SDS-PAGE and mass spectrometry (Fig 1-1). Despite reports that the HAT module of NuA4 can also form an independent complex in yeast^16^, we did not observe smaller particles that would correspond in our cryo-EM analysis (Fig 1-2). Using single particle cryo-EM image analysis we confirmed that the NuA4 complex contains a rigid central hub with more flexible elements attached (Fig 1B,C). Focused refinement of the central hub allowed us to generate a density map for this region with an overall resolution of 3.1Å (Fig 1E, Fig 1-2, Fig 1-4, Fig 1-5) and unambiguously identify the subunits Eaf1, Epl1, Swc4, Arp4, Act1 and the large Tra1 (Fig 1F, Fig 1-5). Missing from this more stable hub are most subunits of the HAT module, all subunits of the TINTIN module, and Yaf9: the three elements known to interact with nucleosomes^31–35^. Our cryo-EM map showed a large diffuse density above the FAT domain of Tra1 (Fig. 1B). While the inherent flexibility and consequently low resolution of these densities does not allow us to determine their subunit composition, evidence suggest that they correspond to the missing HAT and TINTIN modules and Yaf9 subunit.

**Figure 1.**
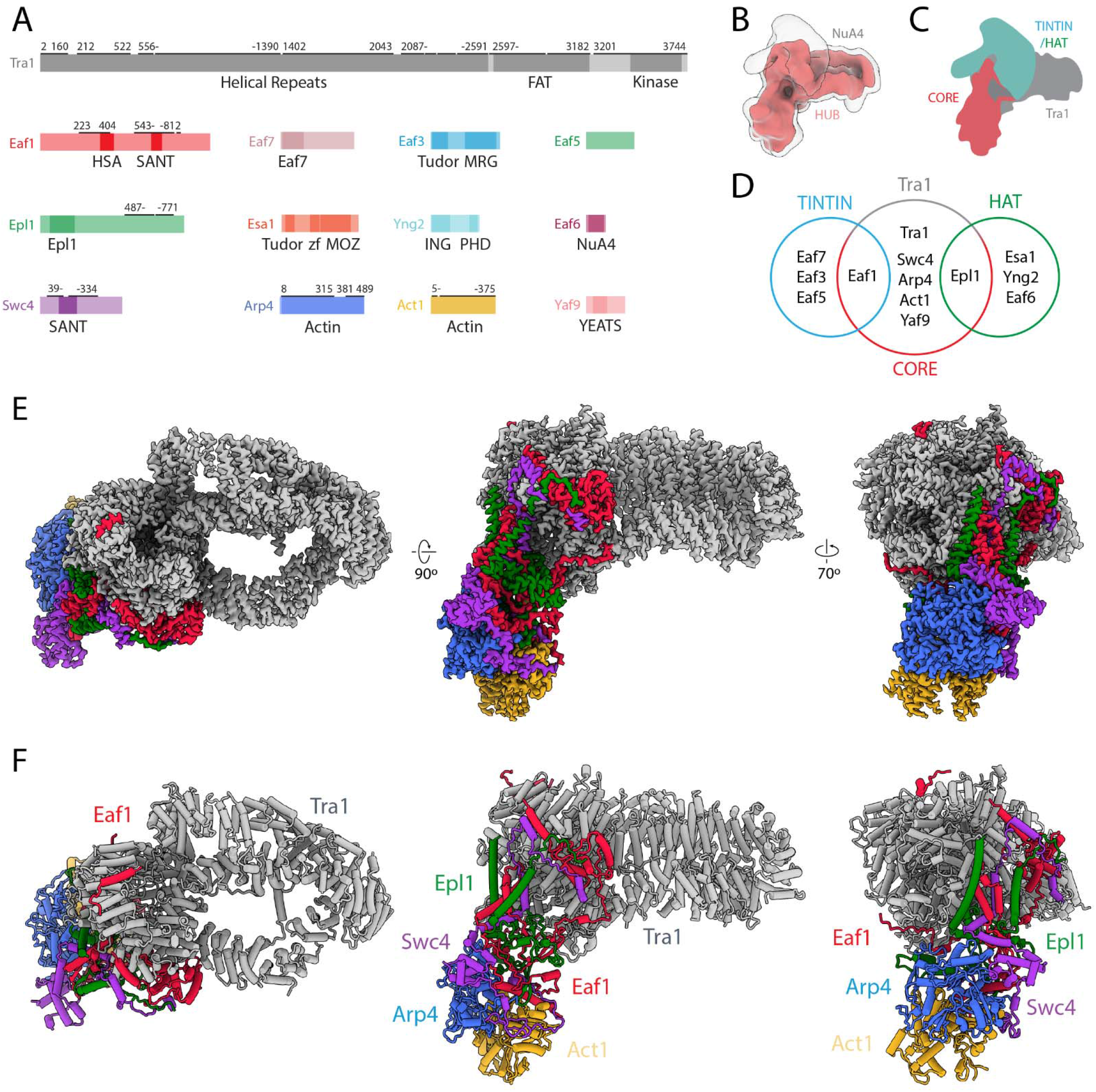
Structure of NuA4. **(A)** Domain map of NuA4 subunits. Modeled regions are marked with a black bar; numbers indicate starting and ending residues. **(B)** Cryo-EM map in red showing the best-defined parts of NuA4. A transparent lower-threshold cryo-EM map is overlaid to show the flexible density likely corresponding to the HAT/TINTIN modules. **(C)** Cartoon representation of NuA4 modules. **(D)** Venn diagram showing NuA4 subunit organization across different complex modules. Subunits in the core attach to Tra1 and act to tether the TINTIN and HAT modules to the complex. **(E)** Cryo-EM map of the NuA4 hub with individual subunits colored. **(F)** Model of the NuA4 hub with individual subunits colored and labelled.

**Figure 1-1.**
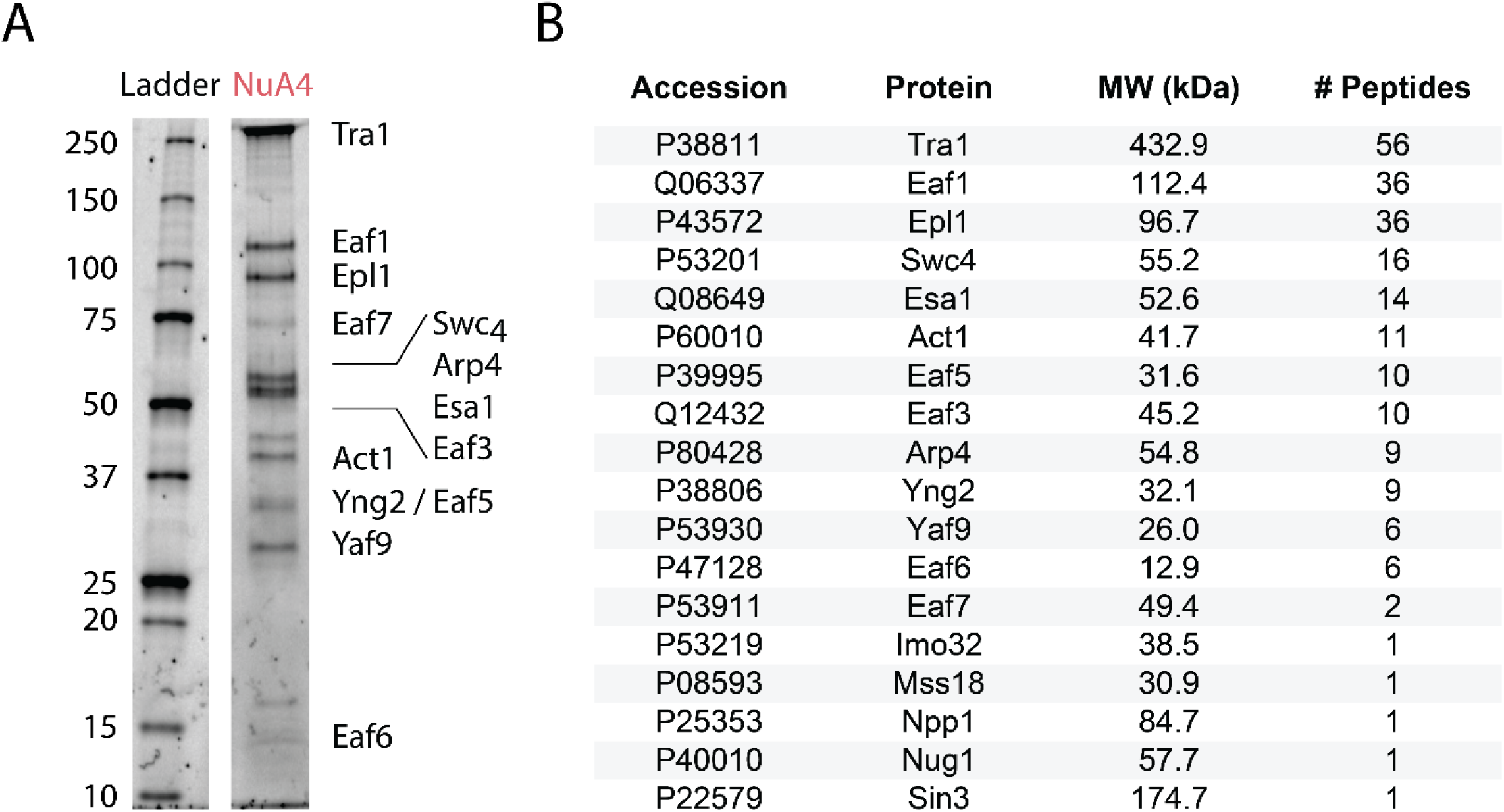
Purification of NuA4. **(A)** SDS-PAGE (BioRad 4–20%) of purified *S. cerevisiae* NuA4, stained with Flamingo (BioRad). **(B)** Mass spectrometry analysis showing the presence of all NuA4 subunits in purified sample.

**Figure 1-2.**
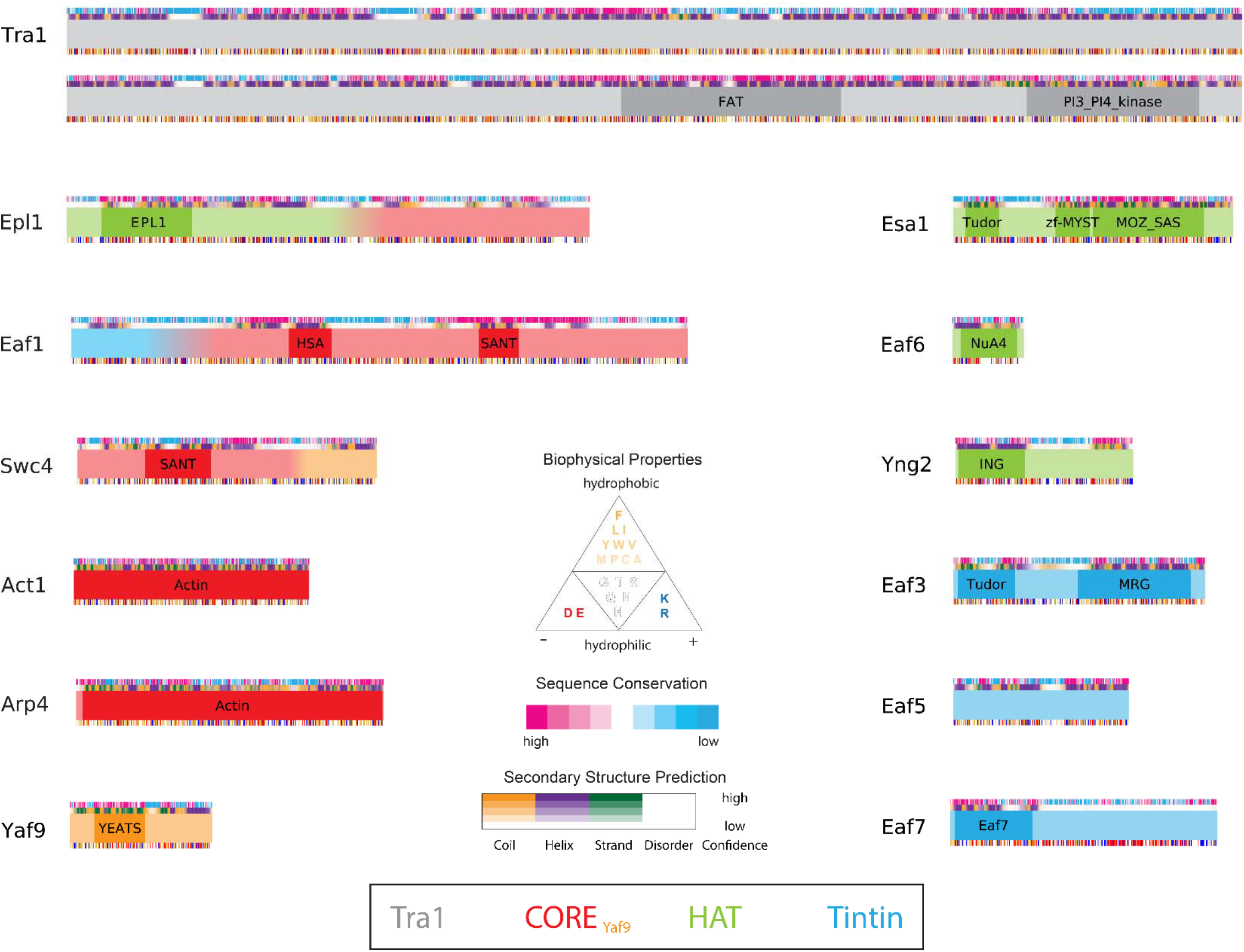
Protein domains of NuA4. Domain maps of NuA4 subunits. Conservation scores ranging from high to low (pink to blue) were generated using the Consurf server^92,93^. PFAM-predicted domains are labeled in bold^107^. Predicted secondary structure based on PSIPRED and DISOPRED3 results (purple for helix, green for stand, orange for coil and white for disordered)^94–96^.

**Figure 1-3.**
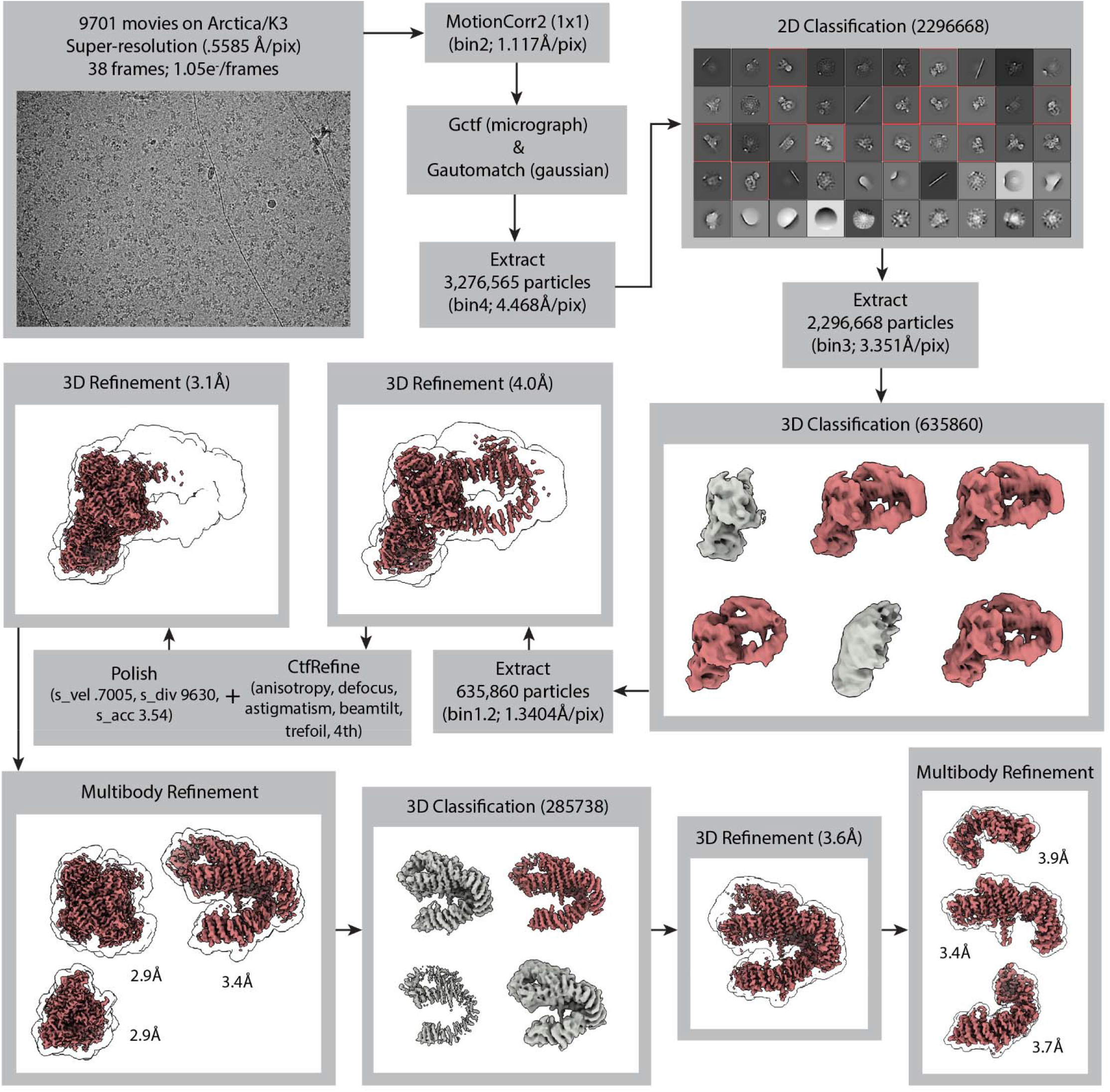
Cryo-EM data collection and processing for NuA4. Cryo-EM data collection and processing for NuA4. Particles from class averages outlined in red and 3D classes colored red were selected for further processing. Particles that went into grey classes were eliminated from further processing. The final global refinement yielded a map with an overall resolution of 3.07Å. To improve map quality in the flexible region of Tra1, multibody refinement was performed for three sections, followed by 3D classification of the partially signal-subtracted particles. The lobes of NuA4, termed core, Tra1-FATKIN and Tra1-HEAT were refined to resolutions 2.9 Å, 2.9 Å and 3.4 Å, respectively. The regions of Tra1-HEAT, termed top, middle and bottom were refined to resolutions 3.9 Å, 3.4 Å and 3.7 Å, respectively.

**Figure 1-4.**
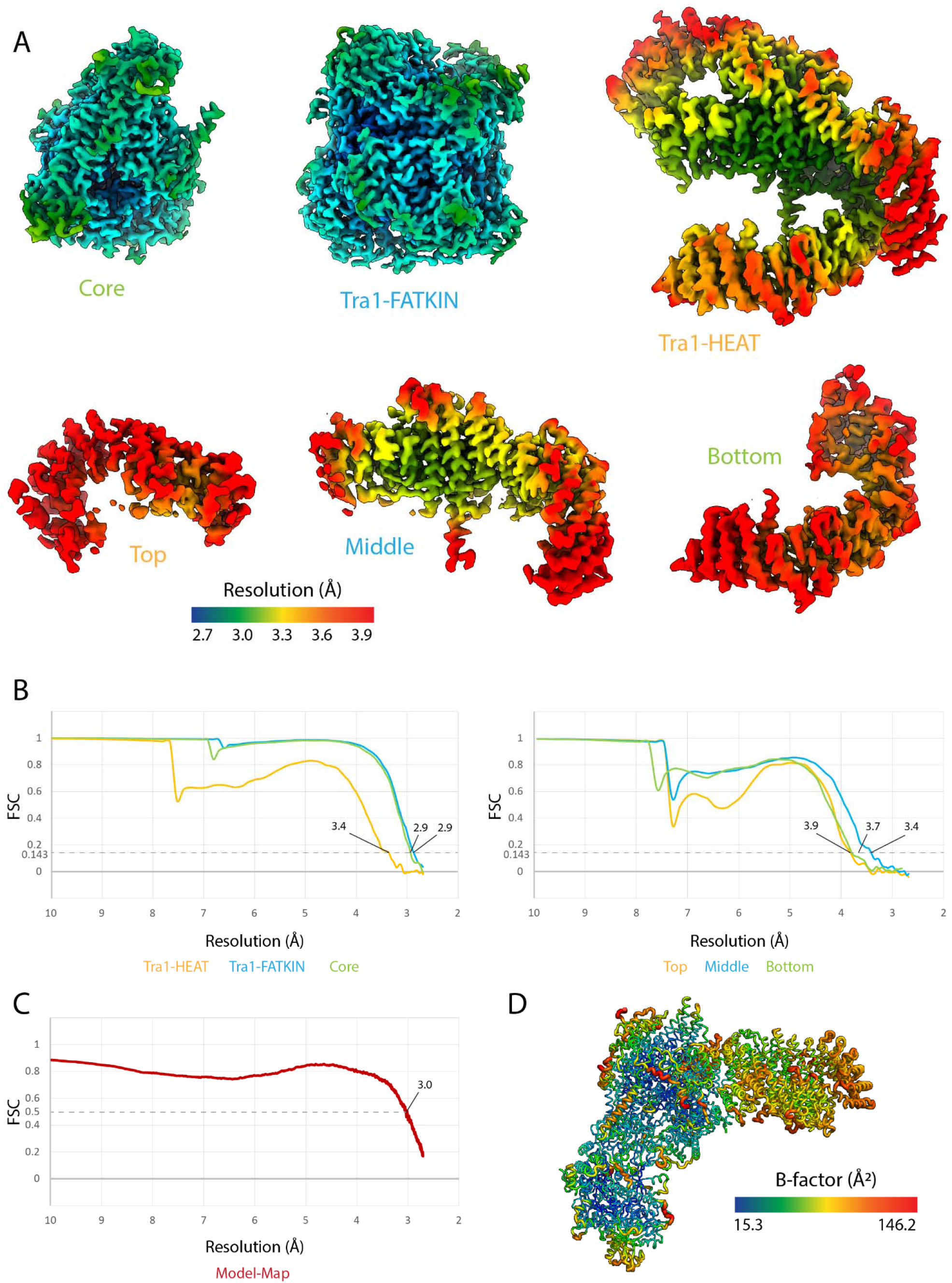
NuA4 structure model validation. **(A)** Multibody maps colored by local resolution^85^. Maps in the top row are from multibody refinement of NuA4. Maps in the bottom row are from multibody refinement of the Tra1-HEAT region. **(B)** Half-map FSC curves of the NuA4 multibody regions (left) and Tra1-HEAT multibody regions (right)^101^. **(C)** Model-map FSC curve (map generated from a composite of multibody maps: core, Tra1-FATKIN, Tra1-HEAT). **(D)** Model of the core of RSC colored according to refined B-factors.

**Figure 1-5.**
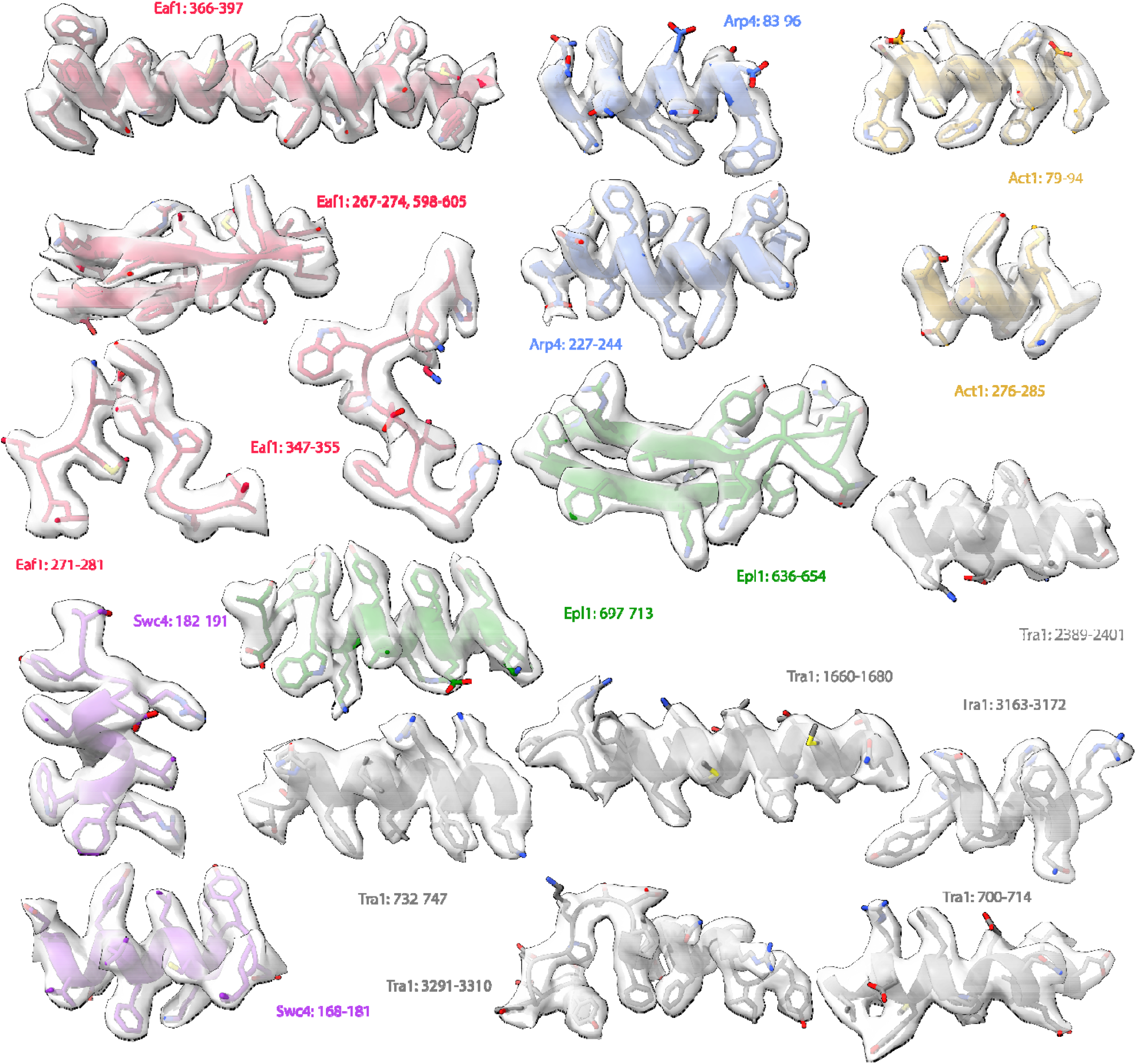
Model fit for NuA4. Transparent regions of the consensus map of NuA4, segmented, with built atomic models shown.

Our cryo-EM based structural model of the central hub was validated by mapping the previously reported chemical crosslinking and mass spectrometry (CX-MS) data of NuA4^30^, which identified many crosslinks between subunits resolved in our structure of the central hub. Overlaying these crosslinks on our structure shows that all 82 identified interactions (52 intra-molecular links and 30 inter-molecular links) fall under the 30Å cutoff for DSS crosslinks (Fig 1-6). This was not the case for the structure of NuA4 previously reported (Fig 1-6)^28^.

**Figure 1-6.**
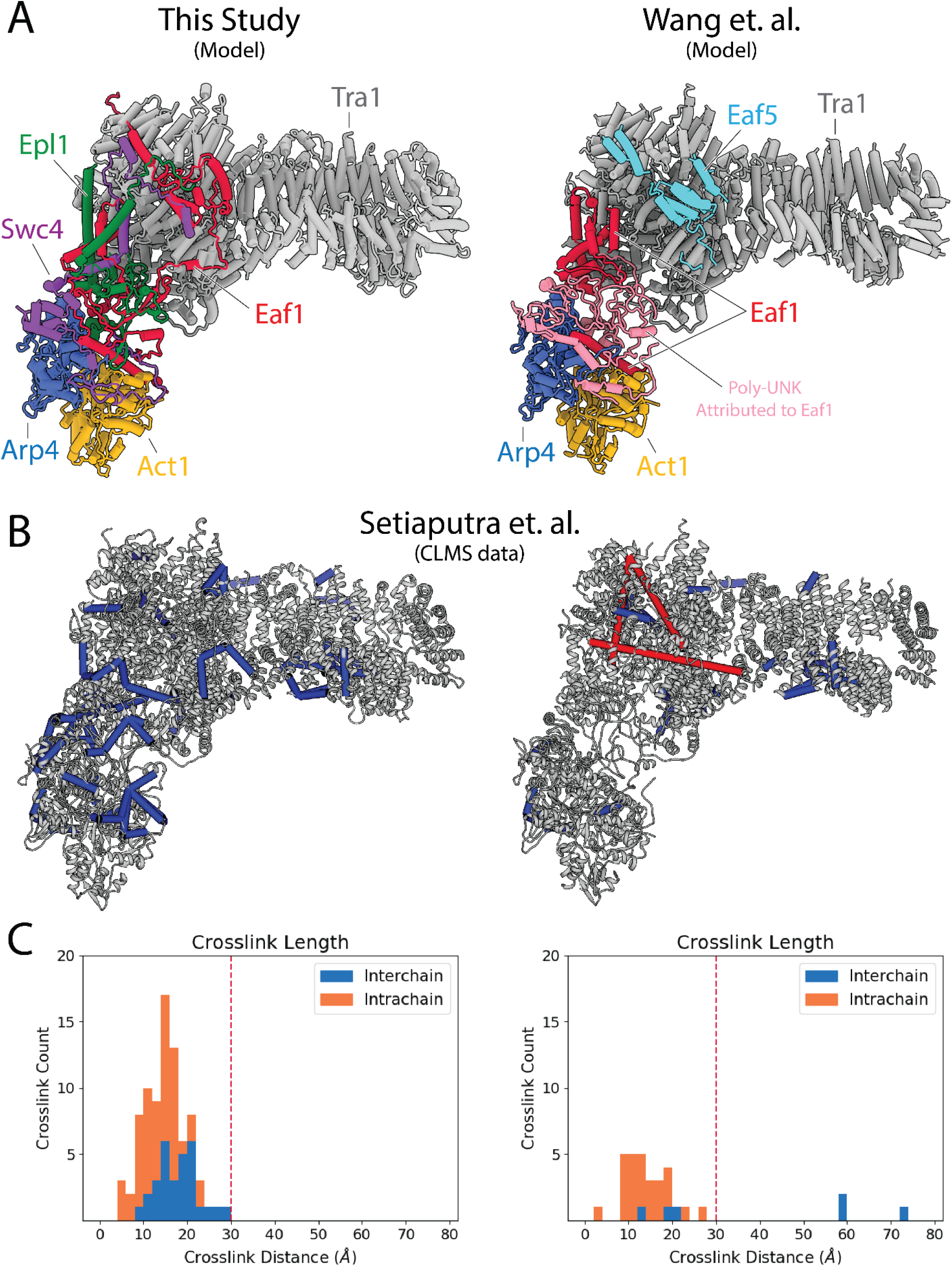
Comparison with previous NuA4 model and CX-MS validation. **(A)** Comparison between the NuA4 model in this study (left) and that of Wang et. al. (right)^28^. In contrast to the previous model, we identify Swc4 and Epl1 within the rigid hub, while do not assign any density to Eaf5. **(B)** Cross-linking mass spectrometry (CX-MS) data of NuA4^30^ mapped on the two models. Crosslinks with a distance >30 Å between alpha carbons are colored red, those with a distance <30 Å are colored green^106^. **(C)** Graph displaying the distribution of crosslink distances between the two models^104^. Inter-chain crosslinks are shown in blue, intra-chain crosslinks in orange. A vertical line indicates a cutoff at 30 Å, the distance considered reasonable for DSS crosslinks.

### Structure of the NuA4 central hub

The structure of the central hub of NuA4 includes Tra1 and a core of additional subunits that interact extensively with each other and tether all the rest of the components (Fig. 1, 2A). The Tra1 subunit makes up most of the hub density (grey in Fig. 1F), and has a very similar structure to that previously described^36^. It contains a large HEAT repeat (pink in Fig. 2), followed by the FAT (yellow in Fig. 2A) and pseudo-kinase domains (jointly also referred to as FATKIN) (cyan in Fig. 2A). Eaf1, Epl1 and Swc4 within the core (red, green, purple, respectively, Fig. 1, 2A) interact with Tra1 near the FATKIN region. Of these, Eaf1 is the primary Tra1 interaction partner, contributing 5,500Å^2^ of the total 7,700Å^2^ buried surface area between the core and Tra1 (Fig. 2C). Arp4 and Act1 (orange and blue, respectively, Fig. 1, 2A) are the only components of the core that do not contact Tra1 (Fig. 2C).

**Figure 2.**
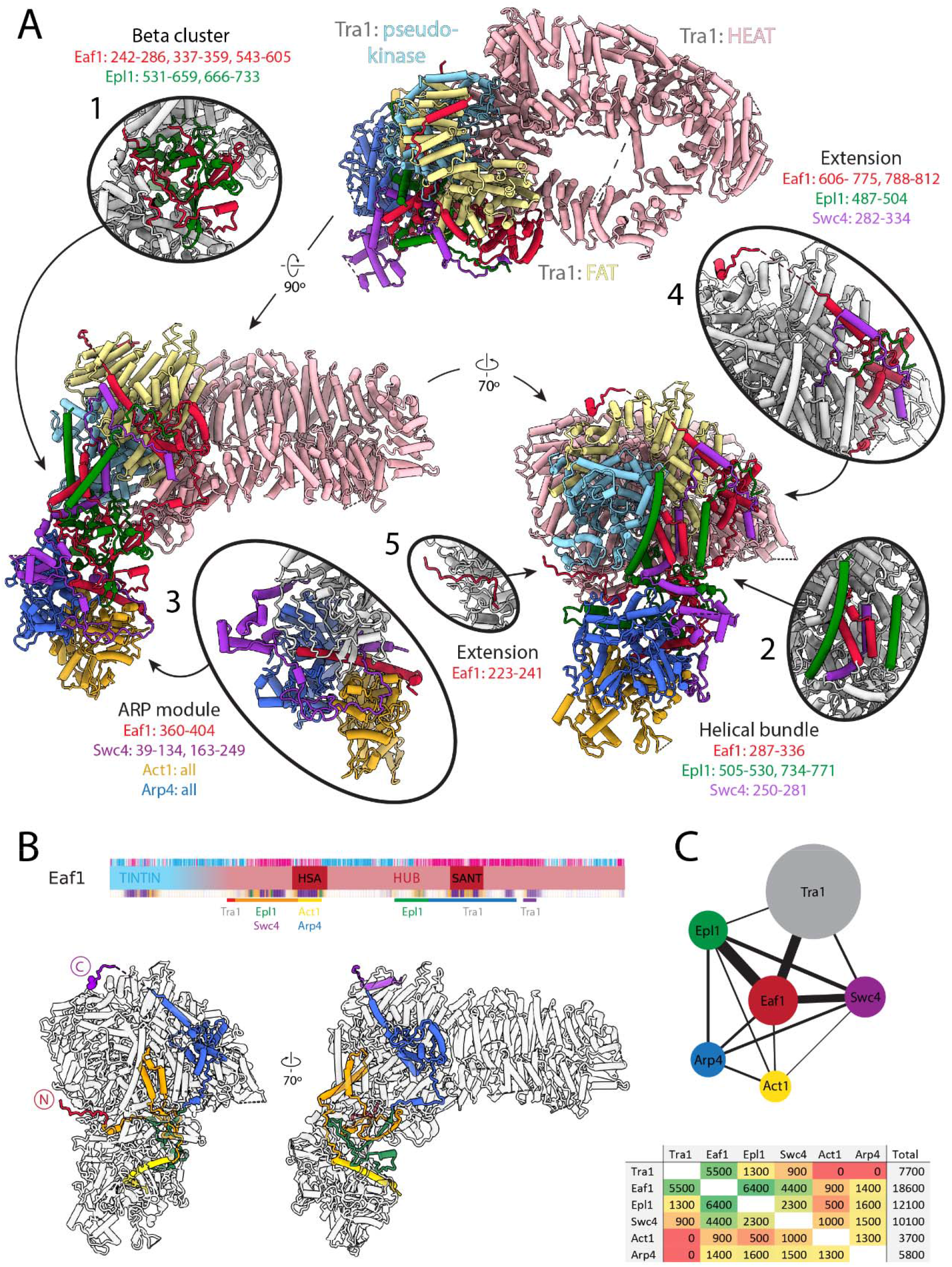
Architecture of the NuA4 central hub. **(A)** Structure of NuA4 with subunits of the core and Tra1 domains in different colors. Within Tra1, the pseudo-kinase domain is colored light blue, the FAT domain is colored pale yellow, and the HEAT domain is colored pink. The FAT and pseudo-kinase domains form the bulk of the interactions with the NuA4 core, highlighted in subpanels (1, 2, 4 and 5). Organization of the NuA4 core can be seen in subpanels (1, 2, 3). Arp module containing Arp4 and Act1 assemble onto the HSA helix of Eaf1 (seen in subpanel 3). **(B)** Top: Eaf1 domain map. Bars underneath are colored to indicate its protein interactions. Bottom: Depiction of Eaf1 interaction with NuA4 subunits. Sections of Eaf1 are colored in a rainbow from N to C terminus with different colors representing regions with different protein-protein interactions. **(C)** Top: Schematic representation of contacts between NuA4 subunits. The width of each line illustrates to extend of the contact area between subunits. Bottom: Table showing the contact area (Å^2^) between NuA4 subunits, colored from red (minimal contact) to green (maximal contact).

Structurally, the NuA4 core made of Eaf1, Epl1, Swc4, Arp4 and Act1, can be seen as containing a beta-cluster, a helical bundle, an Arp module, and two Eaf1 extensions (Fig. 2). The beta cluster is composed of β-strands from Eaf1 and Epl1 (Fig. 2A, oval 1) and sits at the center of the core, surrounded by the helical bundle, Arp module and the Tra1 FAT domain. The helical bundle contains helices from Eaf1, Epl1 and Swc4 (Fig. 2A, oval 2) and buttresses the interface of the Tra1 FAT and pseudo kinase domains. The largest part of the core is the ARP module, which is made up of Act1, Arp4, the HSA helix of Eaf1, and Swc4 (Fig. 2A, oval 3). Act1 and Arp4 bind to each other, end to end, to form a tight dimer. The Swc4 ring, which includes the SANT domain and a long-extended coil, wraps around this Act1/Arp4 dimer, while the long HSA (helicase-SANT–associated) helix of Eaf1 binds across the Arp4/Act1 dimer (similar to other HSA-Arp-actin interactions)^37^. Finally, two sets of Eaf1 extensions emanate from the helical bundle and the beta cluster. The Eaf1 extension from the helical bundle (Fig. 2A, oval 4) binds the FAT domain of Tra1 using its SANT domain and two sets of latch helices. The Eaf1 extension from the beta cluster (Fig. 2A, oval 5) binds the pseudo kinase domain of Tra1 through an extended coil structure.

While most of the components that make up the central hub of NuA4 interact with one another, Eaf1 stands out as the primary scaffolding factor, as it has the largest surface interface with Tra1, Swc4 and Epl1 (Fig. 2B,C). It also contributes the conserved HSA helix to the Arp module. The role of Eaf1 as a scaffolding subunit within NuA4 is in good agreement with previous genetic and biochemical studies that show that its deletion results in loss of NuA4 complex assembly^38,39^.

#### The central hub tethers the nucleosome-interacting components of NuA4

Our cryo-EM reconstruction resolved the structure of the rigid hub of NuA4 containing the core and Tra1, but the three chromatin-interacting components – the HAT and TINTIN modules and the Yaf9 subunit^40,41^ – are missing. These missing parts are flexibly tethered to the core through Epl1, Eaf1 and Swc4, respectively. The missing components likely make up the diffuse density observed above the Tra1 FATKIN domain in our cryo-EM map, as all three have been shown to crosslink with the FATKIN domain (Fig. 3A).

**Figure 3.**
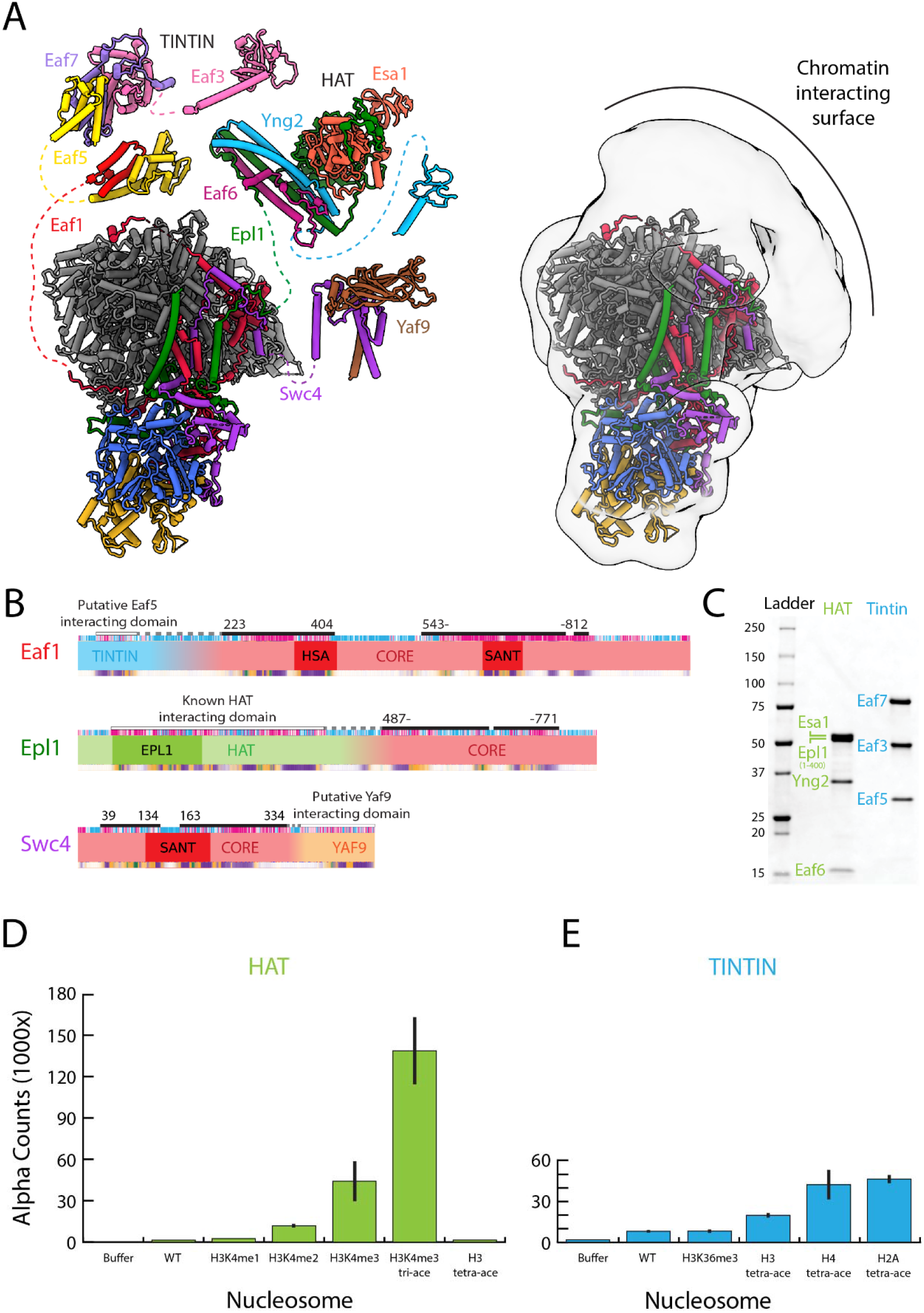
The NuA4 central hub tethers the nucleosome-interacting modules. **(A)** Model for NuA4 complex organization (NuA4 central hub model from the present cryo-EM structure, with the rest of the models from AlphaFold2 prediction)^42,43^. Spatial constraints are imposed on the position of flexible modules by the length of the linker to the connecting aminoacids resolved in the structure. Additional low-resolution density adjacent to the NuA4 hub suggests the appromixate location of the flexible modules. **(B)** Domain map showcasing the subunits that link the TINTIN, HAT and YAF9 modules to the HUB. **(C)** SDS-PAGE (BioRad 4–20%) of purified NuA4 TINTIN and HAT modules, stained with InstantBlue (Expedeon). **(D)** dCypher assay results of nucleosome discovery screen for the purified HAT module. **(E)** dCypher assay result of nucleosome discovery screen for the purified TINTIN module.

Reconstitution experiments and the crystal structure of the HAT module show that it is composed of Yng2, Eaf6 and Esa1, as well as residues 121 to 400 of Epl1^33^. This segment of Epl1 is connected to the rest of the protein in the core of NuA4 via a poorly conserved and predicted unstructured region of about 90 amino acids (residues 487-771) (Fig. 3B), in agreement with the lack of a fixed position for the HAT module with respect to the core. The N-terminus of the region of Epl1 integrated into the core is located near the FAT domain of Tra1, making the HAT module a likely candidate for the diffuse density we see in our structure above the FAT domain (Fig. 3C).

There is little structural information concerning the yeast TINTIN module, which is composed of Eaf3, Eaf5 and Eaf7 and tethered to the core of the complex through Eaf1^30^. We carried out AlphaFold2 prediction^42,43^ of the NuA4 TINTIN module. The resulting structure shows the module is split into three main ordered segments: the Eaf3 chromodomain, the Eaf3 MRG/Eaf5 C-term/Eaf7, and the Eaf5 N-term/Eaf1 (Fig. 3A). These three parts appear interconnected through flexible linkers, making the TINTIN module highly extended. This feature is also captured in the CX-MS data^30^, which shows crosslinks within each of these three parts but no crosslinks between them (Fig. 3-1). The main part of the TINTIN module, composed of Eaf3 MRG /Eaf7/Eaf5 C-term holds the three components of the module together. The interactions between the Eaf3 MRG and Eaf7 are structurally similar to those in the human homolog^44^. The TINTIN module is connected to the NuA4 core through the interaction of the Eaf5 N-term and Eaf1 N-term (aa 28-91). The region of Eaf1 that interacts with the TINTIN module is separated from the Eaf1 region located within the core of NuA4 by a segment of approximately 130 amino acids that is predicted to be unstructured (Fig. 3B). The most N-terminal part of Eaf1 integrated within the core of NuA4 is located near the back of Tra1, near the pseudo-kinase domain, far from the diffuse density seen above the FATKIN domain (Fig. 3A). However, the long linker length between the integrated region of Eaf1 and the region that is predicted to interact with the TINTIN module could easily span the distance to the diffuse density and enable the observed crosslinks between Eaf5 and Tra1 to form. So, in addition to the HAT domain, the TINTIN domain could also make up part of the diffuse density seen above the FAT domain in our cryo-EM structure (Fig. 3C).

**Figure 3-1.**
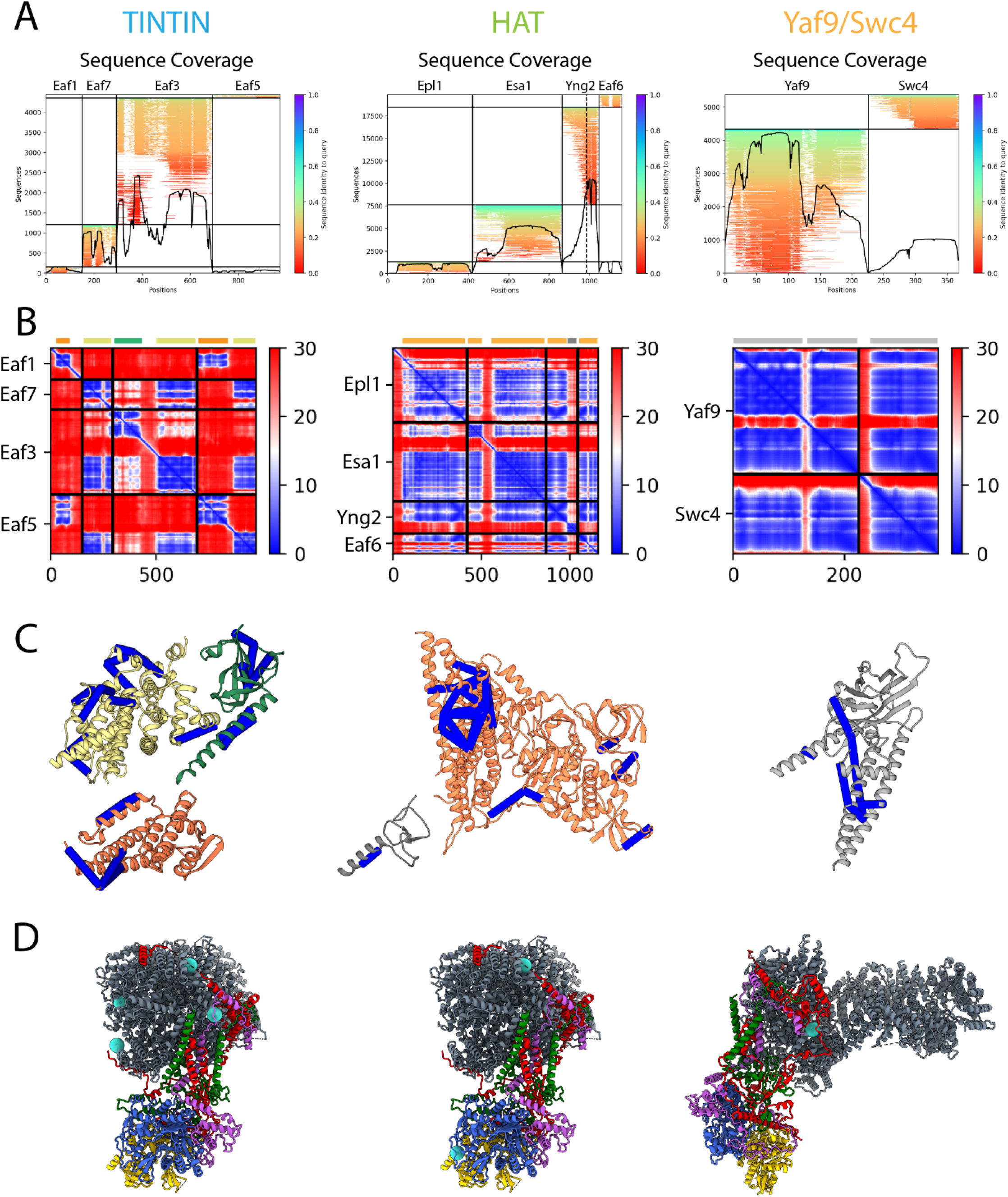
AlphaFold2 prediction and validation of the flexible NuA4 modules. AlphaFold2 prediction for the module structures of TINTIN (left), HAT (middle) and Yaf8/Swc4 (right)^42,43^. **(A)** MSA coverage (MSA search and alignment using Jackhammer) **(B)** Predicted alignment error (blue high, red low) **(C)** AlphaFold2 models (ordered regions) with crosslinks mapped (blue <30 Å, red >30 Å)^30,106^. **(D)** NuA4 crosslinking position with the given module indicated by cyan spheres^30^.

**Figure 3-2.**
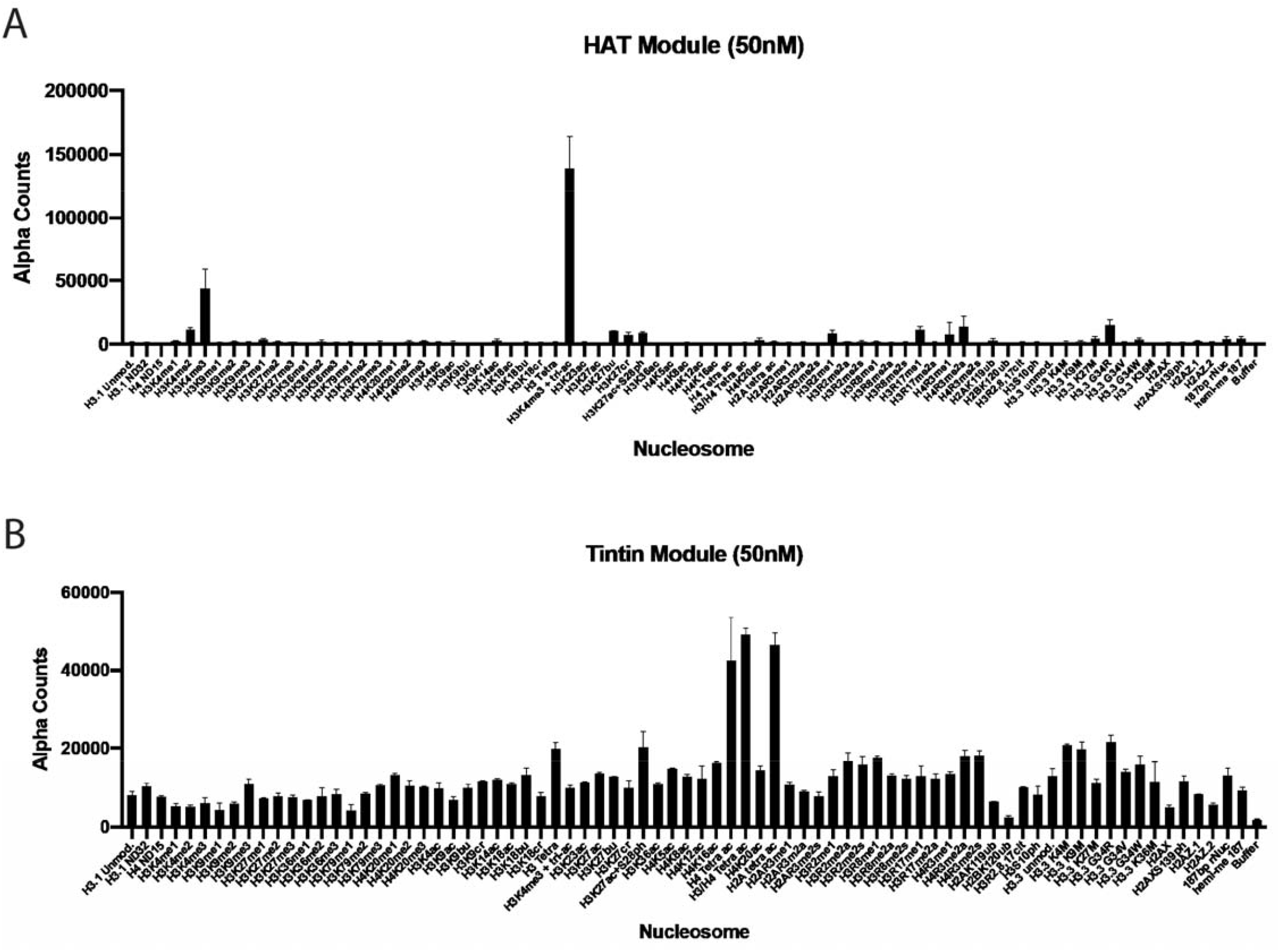
Complete results of dCypher nucleosome discovery screen. **(A)** dCypher nucleosome discovery screen binding profile for purified HAT module. **(B)** dCypher nucleosome discovery screen binding profile for purified TINTIN module.

**Figure 3-3.**
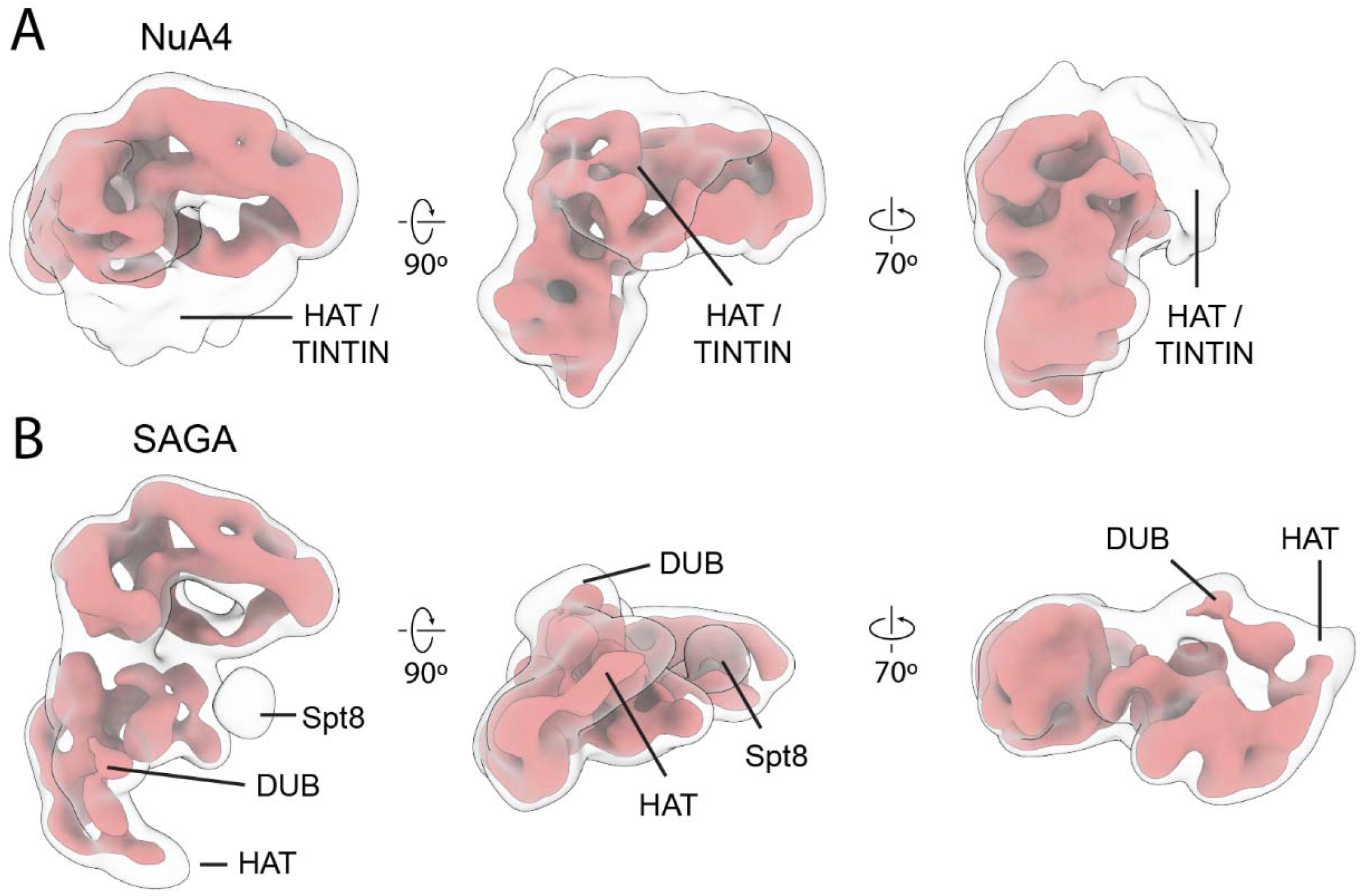
NuA4 and SAGA both have flexible nucleosome engaging domains. Negative stain reconstructions of NuA4 **(A)** and SAGA **(B)** at two thresholds: high (solid red) and low (transparent white). Low threshold shows the flexible chromatin interacting modules of the two complexes. SAGA domain assignments based on previous structural studies^45,76,77^.

**Figure 3-4.**
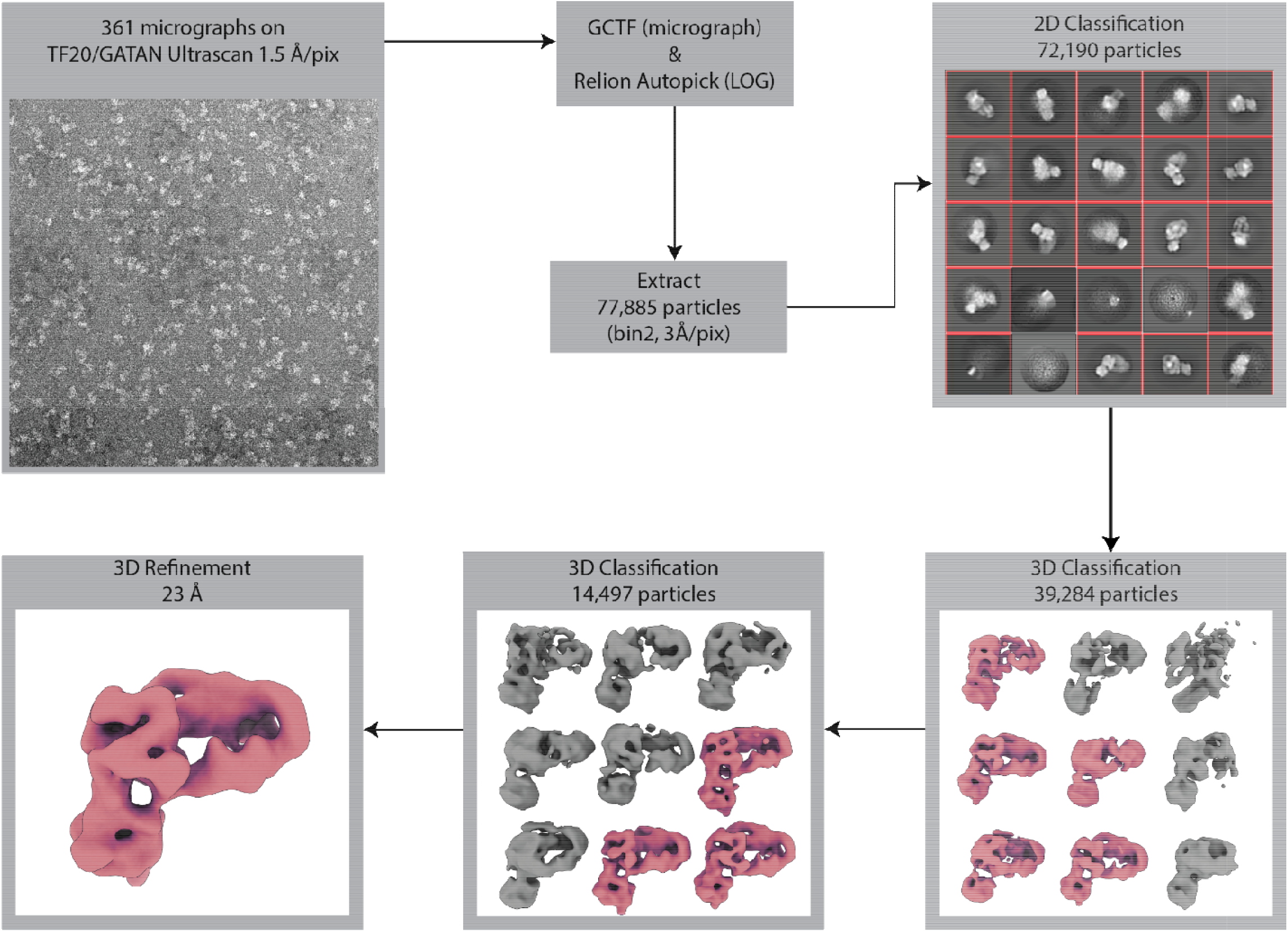
NuA4 NS processing. Negative stain EM data collection and processing for NuA4. Collected images were subjected to CTF estimation using GCTF and the picked and analyzed by 2D and 3D classification in Relion^84,85^.

**Figure 3-5.**
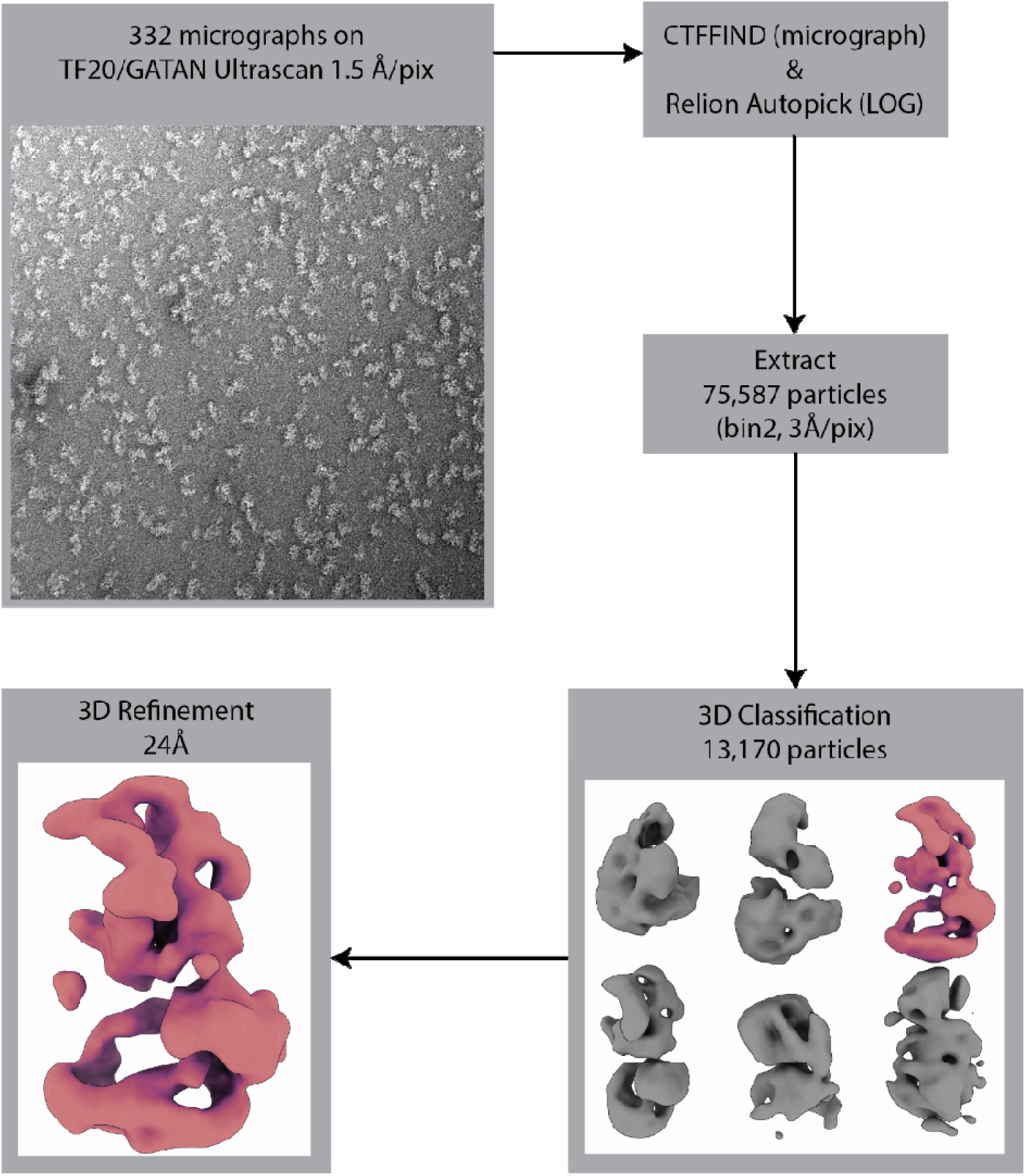
SAGA NS processing. Negative stain EM data collection and processing for SAGA. Collected images were subjected to CTF estimation using GCTF and the picked and analyzed by 3D classification in Relion^84,85^.

Finally, in NuA4 (and the SWR remodeling complex), CX-MS data show that Yaf9 interacts with the C-terminus of Swc4 (residues 359-476)^30^. Previous binding studies have shown that the YEATs domain of Yaf9 is capable of binding acetylated lysine residues, with the highest affinity for H3K27ac^41^. Such interaction has been visualized by an X-ray crystal structure of the Yaf9 YEATS domain bound to an acetylated peptide^41^. The AlphaFold2 prediction of Yaf9 and the C-terminus of Swc4 shows that the Swc4 binds the Yaf9 β-sandwich on the opposite side of the histone binding face and forms a coil-coil structure with a protruding C-terminal helix on Yaf9 (Fig. 3A, Fig. 3-1). The region of Swc4 that is integrated into the core of NuA4 and interacts with Yaf9 is linked via a poorly conserved and predicted unstructured region of about 20 amino acids (Fig. 3B). So, like the HAT and TINTIN module the Yaf9 could also occupy the diffuse density above the FAT domain of Tra1.

NuA4 is not the only HAT to have its various chromatin-interacting and modifying modules flexibly tethered to the core of the complex. Our negative stain electron microscopy visualization of NuA4 and yeast SAGA show how the two complexes similarly contain flexible elements that give rise to poorly defined density (Fig. 3-3A,B,3,4), which in SAGA have been previously mapped to correspond to the HAT and DUB modules^45^. Thus, the NuA4 and SAGA complexes, which share the Tra1 subunit that interacts with site-specific transcription factors, also have a similar overall architecture that includes flexible attachment of chromatin binding and modifying modules. We propose that such organization of functional modules likely reflects a similar chromatin targeting mechanism for these two large acetyltransferase complexes.

#### The HAT and TINTIN modules recognize active transcriptional marks

Due to the small amounts of NuA4 that can be purified from yeast and the generally weak chromatin binding activity of the complex, testing the chromatin recognition capabilities of NuA4 is difficult^46^. To overcome these constraints, we reconstituted the HAT and TINTIN modules separately through recombinant expression and utilized the dCypher approach to interrogate the binding of the two modules against PTM-defined histone peptides (data not shown) and nucleosomes (Fig. 3C)^47–49^.

The HAT module contains three putative chromatin interacting domains: a chromodomain and the HAT domain in Esa1 (the later combines a zinc-finger and MOZ-type HAT domain), and a PHD in Yng2. Previous reports have shown that the PHD domain of Yng2 can bind H3K4me3, while the HAT domain binds the histone octamer surface of the nucleosome^33,40^. Although we did see increasing HAT affinity for nucleosomes with a greater number of methyl groups on H3K4, affinity for nucleosomes containing both H3K4me3 and acetylation marks on K9, K14 and K18 demonstrated ∼3x binding preference over H3K4me3 alone (Fig. 3D, Fig. 3-2A). The tetra-acetylated H3 nucleosome (K4, K9, K14 and K18), however, was not bound by the HAT module, indicating that the acetylation itself is not recognized (Fig. 3D). Instead, the acetylated lysines likely reduce overall interactions between the histone tails and DNA, allowing the tails greater flexibility and the HAT module greater access to the H3K4me3 mark^9,49–51^.

Within the TINTIN module, the only predicted chromatin interacting domain is the Eaf3 chromodomain. Eaf3 is also present in the yeast HDAC RPD3S, where it has been proposed to recognize H3K36me3 modified nucleosomes^52–55^. However, we found that TINTIN has overall weak affinity for nucleosomes and no specificity for H3K36me3 (Fig. 3E, Fig. 3-2B)^46,56^. The only slight preference of the complex appears to be for H2A and H4 acetylated nucleosomes, though with the caveat that this was observed under conditions of high binding background (Fig. 3E, Fig. 3-2B). Interestingly however, these two sets of histone modifications are made by the HAT module of NuA4.

Our binding studies suggest that NuA4 binding is enhanced towards H3K4me3 containing nucleosomes that are hyperacetylated on H3. Acetylation of H3 is largely controlled by SAGA in yeast, in the context of transcription and double strand break repair^11,57^. This suggests that NuA4 activity may follow SAGA activity but precede the recruitment of factors like TFIID or SWR1, which are capable of sensing H4 acetylation^58–60^.

#### Structural comparison of NuA4 with other complexes

Many of the components of yeast NuA4 are shared with other complexes^21,24^. Part of the ARP module is also found in the SWR1 and INO80 chromatin remodelers (Fig. 4)^26,61^, while the large activator targeting Tra1 subunit is shared with SAGA, the other major HAT with functions in transcription^24^. The HAT and TINTIN modules have also been observed distinct from NuA4^16,62^. Of note, the higher eukaryotic TIP60 complex combines components from the yeast NuA4 and SWR1 complexes^38^.

**Figure 4.**
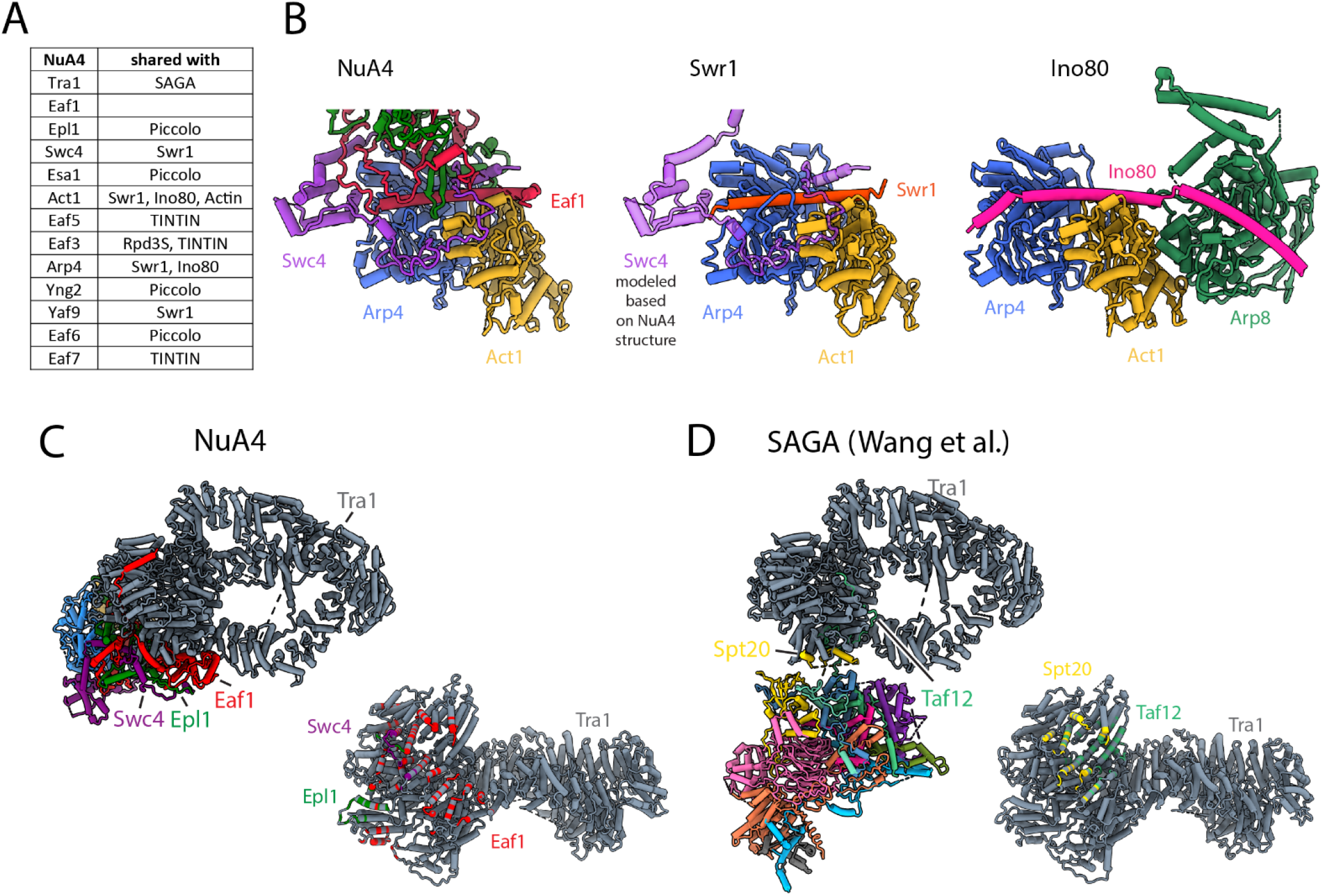
Subunit sharing. **(A)** Table showing protein subunits of NuA4 and the complexes in yeast that share these subunits. **(B)** Comparison of Arp4-Act1 interactions with HSA helix in NuA4 (Eaf1 subunit), SWR1 (Swr1 subunit) and INO80 (Ino80 subunit)^70,71^. **(C)** Left/Top: Model of NuA4 depicting attachment of the core to Tra1. Right/Bottom: The Tra1 subunit of NuA4 colored according to which protein chain contacts it. **(D)** Left/Top: Model of SAGA depicting attachment of the core to Tra1. Right: The Tra1 subunit in SAGA colored according to which protein chain contacts it^76,77^.

The yeast INO80 and SWR1 complexes function as histone remodelers / exchangers during transcription initiation and DNA repair^26,61,63,64^. These functions are in part facilitated by NuA4, which acetylates the nucleosomes that are to be remodeled or have their histones exchanged^65–68^. In SWR1, the ARP module contains Act1, Arp4, Yaf9, and the HSA helix of Swr1 replacing Eaf1 (Fig. 4A, B)^69,70^. In INO80 the ARP module contains Act1, Arp4 and the HAS helix of Ino80 (Fig. 4B)^71^. In both SWR1 and INO80, the ARP module is flexible with respect to the core of the complex, with the HSA helix of Swr1 and Ino80 predicted to be solvent exposed^71–75^. From biochemical studies, the solvent exposed side of the HSA helix in INO80 binds extra-nucleosomal DNA, as also predicted for SWR1^71,72^. In NuA4, the ARP module surface with the HSA helix is buried in the core of the complex, where it appears to serve largely a structural role (Fig. 4B).

In *S. cerevisiae* the multi-subunit NuA4 and SAGA acetyltransferase complexes both include the Tra1 transactivation-binding protein^25^. Of note these two complexes do not share any other subunits and acetylate different histone tails (NuA4 targeting H2A / H4; SAGA targeting H3). Despite their compositional and functional differences, both complexes contain a central core that interacts with Tra1, primarily through the FAT domain, and anchors the rest of the functional domains within them. The NuA4 core contacts Tra1 through an extensive interface involving Eaf1, Swc4 and Epl1. In contrast, yeast SAGA has a much smaller Tra1 interface involving Spt20 and Taf12 (Fig. 4C)^76,77^. The larger interface between Tra1 and the NuA4 core results in a more rigid connection between these two modules compared with yeast SAGA (Fig. 4C)^76,77^.

The human TIP60 complex encompasses the functionalities of yeast NuA4 and SWR1 complexes^21,38^, with the metazoan EP400 being the key protein responsible for merging the two complexes present in lower eukaryotes^38^. EP400 likely evolved through the insertion of the ATPase module of Swr1 between the HSA helix and SANT domain of Eaf1 (Fig. 4-1A,B)^38^. We would like to propose a model in which the N-terminus of EP400 would be within the NuA4 portion of the TIP60 complex, forming part of the core and contributing the HSA helix to the ARP module. The next section of EP400 would go into the SWR1 portion of the complex, where it would contribute the Snf2 domain of the ATPase module, the RUVBL interaction domain, and the helicase domain of the ATPase module. From there, the C-terminal segment of EP400 would return to the NuA4 portion of the complex, where it would make further contributions to the core, ending with the interaction of its SANT domain with the TRRAP subunit (Tra1 homolog) (Fig. 4-1C)^73^.

**Figure 4-1.**
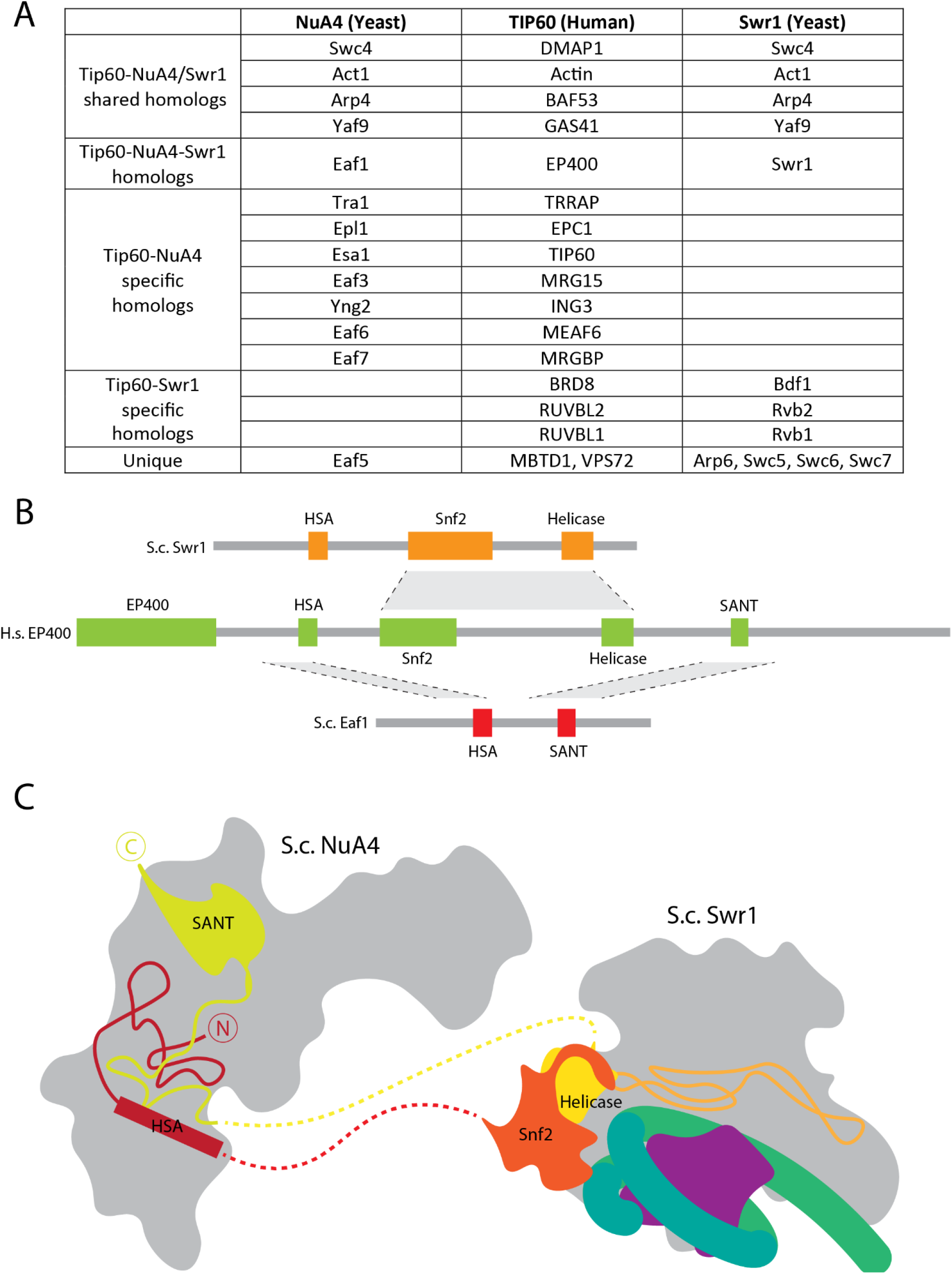
A model of *H*.*s*. TIP60. **(A)**Table of homologous protein subunits in budding yeast (S.c.; Saccharomyces cerevisiae) NuA4, *S*.*c*. Swr1 and human (*H*.*s*.; Homom Sapiens) TIP60. **(B)**Domain similarities between *S*.*c*. Swr1 and *S*.*c*. NuA4 with *H*.*s*. EP400. **(C)**Schematic for the likely structural organization of human TIP60 based on the structures of yeast NuA4 and Swr1^73^.

#### Model of nucleosome selection and acetylation by NuA4

From our structural and biochemical data, we propose a sequential model of NuA4 recruitment to target nucleosomes (Fig. 5). The first step would be the binding of site-specific transcription factors, capable of interaction with the Tra1 subunit to recruit NuA4. The NuA4 flexible HAT and reader modules (TINTIN and Yaf9) would then be able to select neighboring nucleosomes containing preferred modifications, such as H3K4me3 with acetyl groups on H3 K9, K14 and K18. Once a nucleosome is selected by the complex, the HAT module would acetylate histones H2A and H4.

**Figure 5.**
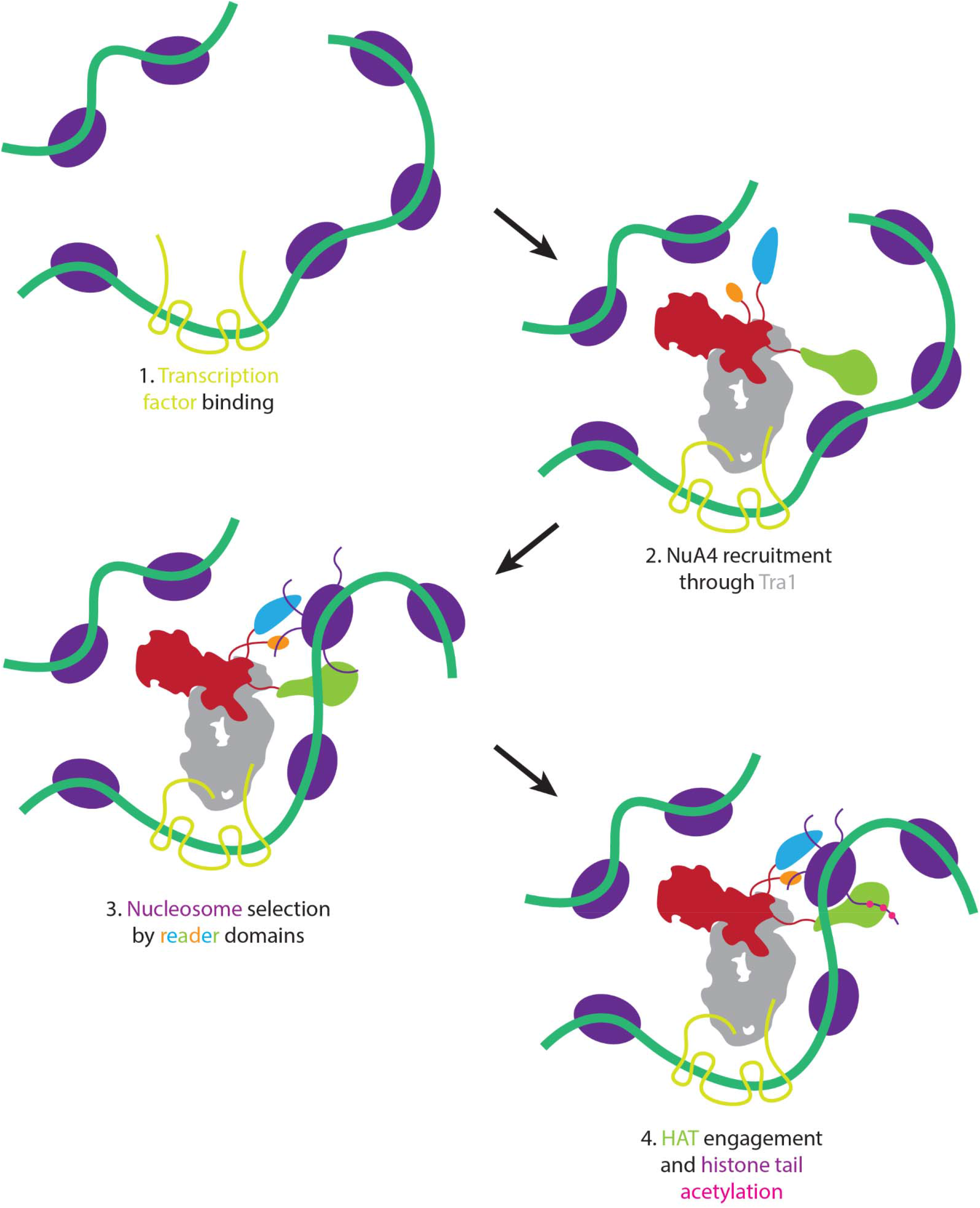
Proposed model of nucleosome selection and propagation of acetylation. Model of NuA4 chromatin localization and histone acetylation. NuA4 is recruited to genomic loci through interaction of Tra1 with site-specific transcription factors. Once recruited, the flexible reader domains interrogate nearby nucleosomes. Nucleosomes bearing the proper chemical marks are preferentially acetylated.

This NuA4 mechanism of recruitment is likely similar for the human TIP60 homolog and may also be shared by other HAT complexes such as SAGA and CBP/p300. Like NuA4, SAGA and CBP/p300 also have their activator-binding and HAT components flexibly separated^76–79^. This structural arrangement likely means that recruitment of the complexes does not constrain the HAT module to act on a specific nucleosome, such as the most adjacent one, but could allow for the complex to target specifically modified nucleosomes within the neighboring chromatin environment.

## Methods

### Protein purification

#### NuA4 and SAGA

NuA4 and SAGA were purified from *Saccharomyces cerevisiae* using a modified TAP purification as described^80^. A strain modified with a TAP tag on the C terminus of ESA1 or SPT7 (*GE Dharmacon*) was grown at 30□C in YPD. 12 L of cells were harvested at OD ∼6.0 and lysed using a Planetary Ball Mill (Retsch PM100). The ground cells were resuspended in lysis buffer (100 mM HEPES pH 7.9, 2 mM MgCl_2_, 300 mM KCl, 10 µM leupeptin, 0.5 mM PMSF, 1X EDTA-free cOmplete protease inhibitor, 0.05% NP40, 10% glycerol) and dounced until homogenous. The lysate was centrifuged for one hour at 15,000 g, and clarified lysate passed through a clean column frit, and aliquoted into 50 mL falcon tubes. To each 50mL tube of lysate, 200 µL of washed IgG resin was added and allowed to incubate overnight at 4°C with gentle rocking. After incubation, resin was collected by centrifuging tubes at 2000 g for 10 minutes and removing the supernatant. Resin was resuspended in 400 µL of TAP buffer (20 mM HEPES pH 7.9, 2 mM MgCl_2_, 300 mM KCl, 10 µM leupeptin, 0.5 mM PMSF, 0.05% NP40, 10% glycerol) and transferred to a microcentrifuge tube. Tubes were centrifuged for 2 minutes at 2000 g and supernatant removed. Resin was washed 5X with TAP buffer. To washed resin, one volume of TAP buffer was added, then TCEP was added to a concentration of 1 mM. 40 µg of TEV protease (Macrolab) was added for every 500 µL of resin, and tubes were let incubate at 4°C overnight.

Tubes were centrifuged for 2 minutes at 2000 g, and the supernatant transferred to another tube as the first elution. Resin was washed 2X with one volume of TAP buffer, and supernatant saved as 2^nd^ and 3^rd^ elutions. All elutions were run on SDS-PAGE and flamingo stained. Those containing NuA4 complex (based on SDS-PAGE and later confirmed by mass spectrometry) were pooled, and the calcium concentration increased by adding calcium chloride to 2 mM final by fast dilution. Pooled elutions were incubated with 150 uL washed CBP resin at 4°C for four hours. Tubes were centrifuged for 2 minutes at 2000 g, and the supernatant removed. Resin was washed 5X with one volume of CBP wash buffer (20 mM HEPES pH 7.9, 2 mM MgCl_2_, 300 mM KCl, 2 mM CaCl_2_, 10 µM leupeptin, 0.01% NP40, 10% glycerol). To washed resin, one volume of CBP elution buffer (20 mM HEPES pH 7.9, 2 mM MgCl_2_, 300 mM KCl, 2 mM EGTA, 10 µM leupeptin, 0.01% NP40, 10% glycerol) was added and let incubate for 30 minutes. Tubes were centrifuged for 2 minutes at 2000g, and the supernatant was transferred to another tube. Elution was repeated three times. Elutions were run on SDS-PAGE and flamingo stained, with those containing NuA4 or SAGA flash-frozen in liquid nitrogen.

#### HAT

The NuA4 HAT module was purified from Hi5 insect cells (*Trichoplusia ni*) co-expressing Yng2, Eaf6, Esa1 and Epl1 (1-400). All genes were codon optimized and synthesized as gBlocks (*IDT*), with Epl1 harboring a C-terminal HIS tag. Synthesized genes for Yng2, Eaf6 and Esa1 were cloned into plasmid 438A. while Epl1 (1-400) was cloned into plasmid 38Q (N-terminal MBP - TEV site)^81^. Genes were then combined into a single plasmid^81,82^ used to generate bacmids by transformation into DH10MultiBac cells (Macrolab). Purified bacmids were transfected into Sf9 cells (*Spodoptera frugoperda*) using FuGene (Promega) and baculoviruses amplified for two rounds before protein expression in Hi5 cells.

Harvested cells were resuspended in 50 mL of lysis buffer (25mM HEPES 7.9, 150mM NaCl, 2.5mM MgCl_2_, 10%glycerol, 1X cOmplete EDTA-free protease inhibitor (Roche)) and sonicated for a total processing time of 5 minutes at 40% power. 5µL of benzonase (Sigma) was added and incubated at 4°C for 20 minutes with gentle rocking. NaCl was added to a final concentration of 300 mM, as well as 50µL of 10% Triton-X and 50µL of NP-40 substitute. Lysate was incubated at 4°C for 10 minutes with gentle rocking. Lysate was centrifuged for 50 minutes at 18,000 rpm and supernatant was removed.

Imidazole, to a final concentration of 10 mM was added to supernatant, which was then incubated with 1.5 mL washed and packed nickel resin (GoldBio) at 4°C for one hour with gentle rocking. Beads were removed and washed five times with 10 mL of wash buffer (25mM HEPES 7.9, 300mM NaCl, 2.5mM MgCl_2_, 10% glycerol, 25 mM Imidazole). Protein was eluted with elution buffer (25mM HEPES 7.9, 300mM NaCl, 2.5mM MgCl_2_, 10% glycerol, 250mM Imidazole). Elutions were analyzed with SDS-PAGE, and samples containing NuA4 HAT were pooled. TCEP was added to a concentration of 1mM, and pooled elutions were incubated with 4mL of washed and packed amylose resin (GE) at 4°C for one hour with gentle rocking. Beads were removed and washed five times with 10 mL of amylose wash buffer (25mM HEPES 7.9, 300mM NaCl, 2.5mM MgCl_2_, 10% glycerol). Protein was eluted with amylose elution buffer (25mM HEPES 7.9, 100mM NaCl, 2.5mM MgCl_2_, 10% glycerol, 20 mM Maltose). Fractions containing NuA4 HAT were pooled and concentrated to 9mg/mL. Sample was incubated with TEV protease (Macrolab) overnight at 4°C without any motion. Digested sample was centrifuged at max speed for 10 minutes, then 500µL was loaded onto a 24mL Superdex 200 increase size exclusion column equilibrated into sizing buffer (25mM HEPES, 2.5 mM MgCl_2_, 100 mM NaCl, 10% glycerol). Peaks were analyzed with SDS-PAGE and samples containing the intact NuA4 HAT complex were pooled, concentrated to a final concentration of 5 mg/mL and frozen.

#### TINTIN

The NuA4 TINTIN module was purified from BL21STAR (Macrolab) *E. coli* cells co-expressing Eaf3, Eaf5 and Eaf7 in a polycistronic construct. The polycistronic block was synthesized in two gBlock (IDT) with the individual genes codon optimized and Epl7 harboring a C-terminal FLAG tag. The two gBlocks were cloned together into the 2G-T plasmid (Addgene #29707), so that Eaf3 would have a N-terminal HIS-GST-TEV site^82^. The constructed plasmids were transformed into BL21STAR cells. Cells were grown in TB media at 37°C till they reached an OD of .8, at which point the cells were cooled to 18°C and protein expression was induced with .1mM IPTG. Cells were harvested after 16 hours.

Cells were resuspended in 50 mL of lysis buffer (25mM HEPES 7.9, 150mM NaCl, 2.5mM MgCl_2_, 10% glycerol, 1X cOmplete EDTA-free protease inhibitor) and sonicated for a total processing time of 5 minutes at 40% power. 5µL of benzonase was added and incubated at 4°C for 20 minutes with gentle rocking. NaCl was added to a final concentration of 300 mM, as well as 50µL of 10% Triton-X and 50µL of NP-40 substitute. Lysate was incubated at 4°C for 10 minutes with gentle rocking. Lysate was centrifuged for 50 minutes at 18,000 rpm and supernatant was removed. Imidazole, to a final concentration of 10 mM was added to supernatant, which was then incubated with 1.5 mL washed and packed nickel resin (GoldBio) at 4°C for one hour with gentle rocking. Beads were removed and washed five times with 10 mL of wash buffer (25mM HEPES 7.9, 300mM NaCl, 2.5mM MgCl_2_, 10% glycerol, 25 mM Imidazole). Protein was eluted with elution buffer (25mM HEPES 7.9, 100mM NaCl, 2.5mM MgCl_2_, 10% glycerol, 250mM Imidazole). Elutions were analyzed with SDS-PAGE, and samples containing TINTIN were pooled. Sample was loaded onto a 1mL HiTrap Q column and eluted via salt gradient. Fractions containing TINTIN were pooled and loaded onto a 1mL Heparin column and eluted via salt gradient. Fractions containing TINTIN were pooled, concentrated and loaded onto a 24mL Superdex 200 increase size exclusion column. Peaks were analyzed with SDS-PAGE and samples containing the intact TINTIN complex were pooled, concentrated to a final concentration of 2.5 mg/mL and frozen.

### Negative stain sample preparation and data processing

Purified NuA4 complex was diluted 1:2 with negative stain crosslinking buffer (20 mM HEPES pH 7.9, 0.1mM EDTA, 2 mM MgCl_2_, 1% trehalose, 1% glycerol, 75 mM KCl, 1 mM TCEP, 0.01% glutaraldehyde) and allowed to crosslink on ice for 5 minutes. 4 µL was then applied to a glow-discharged continuous carbon grid for 5 minutes and stained using uranyl formate. The negative stain data set was collected on a Tecnai F20 microscope (FEI) operated at 120 keV and equipped with an Ultrascan 4000 camera (Gatan). Data were collected using Leginon data acquisition software^83^. The CTF parameters were estimated using Gctf (version 1.16) and particles were picked using Relion^84,85^. Data processing was done using Relion (version 3.1)^85^. Extracted particles were subjected to 2D classification, *ab initio* model generation, and 3D classification to identify different conformation states of the complex and to visualize the flexibility of its submodules.

Purified SAGA complex was diluted 1:2 with negative stain crosslinking buffer (20 mM HEPES pH 7.9, 0.1mM EDTA, 2 mM MgCl_2_, 1% trehalose, 1% glycerol, 75 mM KCl, 1 mM TCEP, 0.01% glutaraldehyde) and allowed to crosslink on ice for 5 minutes. 4 µL was then applied to a glow-discharged continuous carbon grid for 5 minutes, stained using uranyl formate, and tilted negative stain data set collected on a Tecnai F20 microscope (FEI) operated at 120 keV / equipped with an Ultrascan 4000 camera (Gatan). Data were collected using Leginon data acquisition software^83^. The CTF parameters were estimated using Gctf (version 1.16), particles picked using Relion^84^., and data processing done using Relion (version 3.1)^85^. Extracted particles were subjected to *ab initio* model generation and 3D classified to identify different conformation states of the complex and visualize the flexibility of its submodules.

### Cryo-EM sample preparation

For cryo-EM sample preparation we used a Vitrobot Mark IV (FEI). NuA4 was crosslinked on ice using 1 mM BS3 (Thermo Fisher Scientific) for 15 min before 4 μL of sample was applied to a graphene oxide coated 1.2/1.3 UltrAuFoil grids (Quantifoil) at 4°C under 100% humidity^86^. The sample was immediately blotted away using Whatman #1 for 2 s at 5 N force and then immediately plunge frozen in liquid ethane cooled by liquid nitrogen.

### Cryo-EM data collection

Grids were clipped and transferred to the autoloader of a Talos Arctica electron microscope (Thermo Fischer Scientific) operating at 200 keV acceleration voltage. Images were recorded with a K3 direct electron detector (Gatan) operating in super-resolution mode at a calibrated magnification of 44,762 (.5585 Å/pixel), using the SerialEM data collection software^87^. 38-frame exposures were taken at 0.065 s/frame, using a dose rate of 12.98 e-/pixel/s (1.05 e-/Å^2^ /frame), corresponding to a total dose of 40 e-/Å^2^ per micrograph (Figure). A total of 9701 movie were collected from a single grid.

### Cryo-EM data processing

All data processing was performed using Relion3 (version 3.0)^85^. Whole movie frames were aligned and binned by 2 (1.117 Å/pixel) with MotionCor2 to correct for specimen motion^88^. The CTF parameters were estimated using Gctf^84^. 3,276,565 particles were picked with Gautomatch (version 0.53, from K. Zhang, MRC-LMB, Cambridge). Particles were extracted binned by 4 (4.468 Å/pixel) and subjected to two-dimensional classification to remove ice, empty picks and graphene oxide creases, which resulted in 2,296,668 particles. These particles were re-extracted binned by 3 (3.351 Å/pixel) and subjected to three-dimensional classification, and the best class, containing 635,860 particles, was selected for further processing. The particles were re-extracted, binned by 1.2 (1.3404 Å/pixel) and subjected to three-dimensional refinement, resulting in a 3.98 Å map. Particles were subjected to CTF refinement and particle polishing before performing another 3D refinement, which resulted in reconstructions at 3.1 Å^85,89^. The core was then subjected to multibody refinement by masking the complex into three parts (core, Tra1-FATKIN, Tra1-HEAT), which refine to 2.9, 2.9 and 3.4 Å^90^. While the core and Tra-FATKIN regions refine to nearly uniform resolution the Tra1-HEAT region showed a wider range of resolutions (and map quality), indicating remaining flexibility within this region. To further refine the Tra1-HEAT region, the core and Tra1-FATKIN regions were subtracted from the particle images and the particle box was re-centered around the Tra1-HEAT region. The resulting particles were subjected to alignment-free three-dimensional classification and a single good class showing the highest resolution features was selected, resulting in 285,738 particles. These particles were refined to 3.6 Å, and subjected to multibody refinement by masking the Tra1-HEAT into three parts (top, middle and bottom – set of helices), which refine to 3.9, 3.4 and 3.7 Å. While the resolution of the Tra1-HEAT region did not improve, the map quality did in parts, allowing better interpretability of the peripheral regions.

Some of the software packages mentioned above were configured by SBgrid^91^.

### Model building and refinement

The multibody maps for the core, Tra1-FATKIN and Tra1-HEAT (top, middle and bottom) were combined for model building. The models for the Tra1 (PDB:5ojs) and Arp module (PDB:5i9e)^36,70^ were rigid-body docked into the combined map and adjusted where needed. The remaining density was manually built using COOT by building poly-alanine chains for all unaccounted density. Each chain segment was then inspected for stretches of high-quality density that could allow for the identification of potential side chain patterns that were then searched for within the sequences of known proteins within the complex (https://github.com/Stefan-Zukin/blobMapper). Chain identification was also aided by secondary structure prediction and sequence conservation^92–96^. The resulting protein model was iteratively refined using PHENIX and manual adjustment in COOT^97,98^. The model was validated using MTRIAGE and MOLPROBITY within PHENIX^99,100^. The refinement statistics are given in Table 1 and show values typical for structures in this resolution range. The FSC curve between the model and the map shows good correlation up to 3.0 Å resolution according to the FSC = 0.5 criterion 101.

**Table 1.** Refinement Table.

| **Data Collection, Map and Model Refinement, Validation** |  |
| --- | --- |
| **Data Collection** |  |
| Dataset | NuA4 |
| Microscope | Talos Arctica |
| Stage type | Autoloader |
| Voltage (kV) | 200 |
| Detector (Mode) | Gatan K3<br>(super-resolution) |
| Pixel size (Å) | 1.117 (.5585) |
| Electron dose (e<sup>-</sup>/Å<sup>2</sup>) | 38 |
| **Reconstruction** |  |
| Software | RELION |
| Particles | 635,860 |
| Box size (pixels) | 320 |
| Map resolution (Å) | 3.1 |
| **Coordinate refinement** |  |
| Software | PHENIX |
| Resolution cutoff (Å) | 3.0 |
| **Model** |  |
| Residues |  |
| Protein | 5259 |
| B-factors |  |
| Protein | 65.28 |
| RMS deviations |  |
| Bond lengths (Å) | .002 |
| Bond angles (°) | .533 |
| **Validation** |  |
| Molprobity Score | 1.75 (87<sup>th</sup>) |
| Molprobity clashscore | 7.36 (85<sup>th</sup>) |
| Rotamer outliers (%) | .02 |
| Cβ deviations (%) | 0 |
| Ramachandran plot |  |
| Favored (%) | 94.93 |
| Allowed (%) | 5.06 |
| Outliers (%) | .02 |

### dCypher binding assays

dCypher binding assays to PTM-defined histone peptides and semi-synthetic nucleosomes were performed as previously^47,48,102^. Briefly, 5 µL of recombinant NuA4 HAT and TINTIN module Queries were titrated against 5 µL of histone peptide (100 nM) or nucleosome (10 nM) Targets and incubated for 30 minutes at room temperature in the appropriate optimized assay buffer (Peptides: 50 mM Tris pH 7.5, 0.01% Tween-20, 0.01% BSA, 0.0004% Poly-L Lysine, 1 mM TCEP; Nucleosomes: 20 mM Tris pH 7.5, [150 mM NaCl for HAT module or 200 mM NaCl for TINTIN module], 0.01% NP-40, 0.01% BSA, 1 mM DTT, [0.4 µg/mL sheared salmon sperm DNA (SalDNA) for HAT module or 0.04 µg/mL SalDNA for TINTIN module]. Then for the 6xHIS-tagged HAT module a 10 µL mixture of 5 µg/mL AlphaLISA Nickel chelate acceptors beads (*PerkinElmer #AL108*) and 10 µg/mL AlphaScreen donor beads (*PerkinElmer #6760002*); or for FLAG-tagged TINTIN module a 10 µL mixture of 1:400 anti-FLAG antibody (*MIlliporeSigma # F7425*), 5 µg/mL AlphaLISA protein-A acceptors beads (*PerkinElmer #AL101*) and 10 µg/mL donor beads were added, followed by a 60 minute incubation. Alpha counts were measured on a *PerkinElmer 2104 EnVision* (680 nm laser excitation, 570 nm emission filter ± 50 nm bandwidth). To optimize salt (NaCl), competitor SalDNA concentrations, and query probing concentration, 2D assays were performed by titrating both query and salt (or SalDNA) against nucleosome substrates (Unmodified (WT), H3K4me3, H3K36me3, and biotin-DNA). It was found that the HAT module had weak DNA binding ability that could be competed away with SalDNA (data not shown). Discovery screens consisting of 77 nucleosome substrates (*EpiCypher #16-9001)* were used to test the HAT (50 nM) and TINTIN (50 nM) modules (Fig. 3-2). All binding interactions were performed in duplicate.

### Creation of figures

Depiction of molecular models were generated using PyMOL (The PyMOL Molecular Graphics System, version 1.8, Schrödinger), the UCSF Chimera package from the Computer Graphics Laboratory, University of California, San Francisco (supported by National Institutes of Health P41 RR-01081) and UCSF ChimeraX developed by the Resource for Biocomputing, Visualization, and Informatics at the University of California, San Francisco, with support from National Institutes of Health R01-GM129325 and the Office of Cyber Infrastructure and Computational Biology, National Institute of Allergy and Infectious Diseases^103–105^. Protein domains graphs (Fig. 1-2, 2, 3) were generated using domainsGraph.py (https://github.com/avibpatel/domainsGraph). Crosslinking mass spectrometry figures were generated using Xlink Analyzer^106^. ColabFold MSA and predicted alignment error figures were generated using ColabFold AlphaFold2-Advanced Notebook (https://colab.research.google.com/github/sokrypton/ColabFold/blob/main/beta/AlphaFold2_advanced.ipynb)^43^.

## Data availability

The cryo-EM maps and coordinate models have been deposited in the Electron Microscopy Data Bank with the accession codes EMD-XXXXX (NuA4 core), EMD-XXXXX (NuA4 Tra1-FATKIN), EMD-XXXXX (Tra1-HEAT), EMD-XXXXX (Tra1-HEAT-top), EMD-XXXXX (Tra1-HEAT-middle), EMD-XXXXX (Tra1-HEAT-bottom) and in the Protein Data Bank with the accession codes PDB-XXXX (NuA4). Plasmids for HAT and TINTIN expression have been made available through Addgene (Catalog #XXXXXX and #XXXXXX).

## Acknowledgements

We thank Anthony Iavarone for mass spectrometry data collection and analysis, Patricia Grob, Daniel Toso and Jonathan Remis for electron microscopy support, Abhiram Chintangal and Paul Tobias for computing support, Basil Greber, Vignesh Kasinath and Michael-Christopher Keogh for feedback on manuscript. This work was funded through NIGMS grant R35-GM127018 to Eva Nogales, and R44GM117683 (AlphaLISA assay development, aka. dCypher) and R44GM116584 (Nucleosome diversity) to EpiCypher. EN is a Howard Hughes Medical Institute Investigator.

## Author contributions

S.A.Z. cloned constructs. S.A.Z. and A.B.P. purified protein, prepared EM samples, collected and processed EM data, and built and refined atomic model. M.R.M. and I.K.P. designed, performed, and analyzed dCypher studies. A.B.P, S.A.Z. and E.N. wrote paper with input from all authors.

